# Soluble MAC is primarily released from MAC-resistant bacteria that potently convert complement component C5

**DOI:** 10.1101/2021.12.15.472789

**Authors:** Dennis J. Doorduijn, Marie V. Lukassen, Marije F.L. van ’t Wout, Vojtech Franc, Maartje Ruyken, Bart W. Bardoel, Albert J.R. Heck, Suzan H. M. Rooijakkers

## Abstract

The Membrane Attack Complex (MAC or C5b-9) is an important effector of the immune system to kill invading microbes. MAC is formed when complement enzymes on the bacterial surface convert complement component C5 into C5b. Although the MAC is a membrane-inserted complex, soluble forms of MAC (sMAC, or terminal complement complex (TCC)) are often detected in sera of patients suffering from infections. Consequently, sMAC has been proposed as a biomarker, but it remains unclear when and how it is formed during infections. Here, we studied mechanisms of MAC formation on bacteria and found that sMAC is primarily formed in human serum by bacteria resistant to MAC-dependent killing. Surprisingly, C5 was converted into C5b more potently by MAC-resistant compared to MAC-sensitive *Escherichia coli* strains. Both the increase in C5 conversion and sMAC generation were linked to the expression of lipopolysaccharide (LPS) O-Antigen in the bacterial outer membrane. In addition, we found that MAC precursors are released from the surface of MAC-resistant bacteria during MAC assembly. Release of MAC precursors from bacteria induced lysis of bystander human erythrocytes in the absence of other serum components. However, serum regulators vitronectin (Vn) and clusterin (Clu) can prevent this bystander lysis. Combining size exclusion chromatography with mass spectrometry profiling, we show that sMAC released from bacteria in serum is a heterogeneous mixture of complexes composed of C5b-8, up to 3 copies of C9 and multiple copies of Vn and Clu. Altogether, our data provide molecular insight into how sMAC is generated during bacterial infections. This fundamental knowledge could form the basis for exploring the use of sMAC as biomarker.

## Introduction

The complement system is a part of the human immune system that plays a crucial role in clearing invading bacteria to prevent infections. The complement system consists of soluble plasma proteins that circulate as inactive precursors [1,2]. When complement is activated at the bacterial surface, a proteolytic cascade is triggered that labels the surface with convertase enzymes [1]. These convertases initially convert C3 into anaphylatoxin C3a and C3b, labelling bacteria for phagocytosis by neutrophils [3]. As terminal step in the pathway, C5 is converted into pro-inflammatory C5a, which recruits and activates neutrophils, and C5b. C5b, together with C6, C7, C8 and up to 18 copies of C9, assembles a large, ring-shaped Membrane Attack Complex (MAC) pore [4–7]. MAC pores can efficiently kill Gram-negative bacteria, although some serum-resistant Gram-negative bacteria can survive killing by MAC pores [8–11], and Gram-positive bacteria are intrinsically resistant to MAC-dependent killing [12]. Nevertheless, the clinical importance of the MAC in humans is evidenced by recurrent infections in patients treated with C5-inhibitor eculizumab [13] or patients with genetic deficiencies in one of the MAC components [14].

Complement activation products that are released into plasma are frequently used as biomarkers for infections [15]. One of these biomarkers is the terminal complement complex (TCC) or soluble MAC (sMAC), which is often increased in plasma of patients suffering from bacterial infections [16–18]. However, since MAC is meant to assemble and insert in bacterial membranes, it is still unclear how sMAC is formed when complement is activated on bacteria, especially in the context of Gram-positive bacteria [17]. sMAC could represent debris of lysed cells, but could also represent improperly inserted MAC pores that are released from bacteria during complement activation [19].

Here, we show that sMAC is primarily formed when complement is activated on bacteria that resist killing by MAC pores. A direct comparison revealed that MAC-resistant *Escherichia coli* strains converted released more sMAC and converted than MAC-sensitive strains. Our data suggest that MAC did not properly insert into the bacterial cell envelope of MAC-resistant strains. Although the release of sMAC could lyse bystander human erythrocytes in a serum-free model, serum regulators vitronectin (Vn) and clusterin (Clu) can prevent this bystander lysis. Finally, combining size exclusion chromatography (SEC) and mass spectrometry (MS) profiling of serum incubated with bacteria revealed that sMAC is a heterogeneous complex composed of C5b, C6, C7, C8, 1 to 3 copies of C9 and several copies of chaperone molecules Vn and Clu. Altogether, our study suggests that sMAC is an inactivated complex released from bacteria that resist killing by MAC.

## Results

### sMAC is primarily formed by MAC-resistant Gram-negative bacteria

To understand how sMAC is formed when bacteria activate complement, we analyzed sMAC formation by different bacteria. A panel of twelve laboratory and clinical *E. coli* strains were incubated with pooled human serum. First, we studied if these *E. coli* strains were sensitive to killing by MAC pores. Bacterial viability was assessed by counting colony forming units (CFU’s) and revealed that four strains were killed in serum (**Fig. 1a**). Killing was MAC-dependent because it could be inhibited with C5 inhibitors OmCI and eculizumab (**S1a**), indicating that these strains are ‘MAC-sensitive.’ The other eight strains survived in serum (**Fig. 1a**). C3 conversion was measured to determine if these serum-resistant strains activated complement at all. Deposition of C3b on the bacterial surface (**Fig. 1b**) and release of C3a in the supernatant (**S1b**) were similar on three strains compared to the MAC-sensitive strains, indicating that these strains activate complement efficiently. We have previously shown that MAC components bind to these strains after complement activation [20], suggesting that they are ‘MAC-resistant’ strains. The other five strains showed little to no deposition of C3b and were considered ‘complement-resistant’.

**Figure 1.**
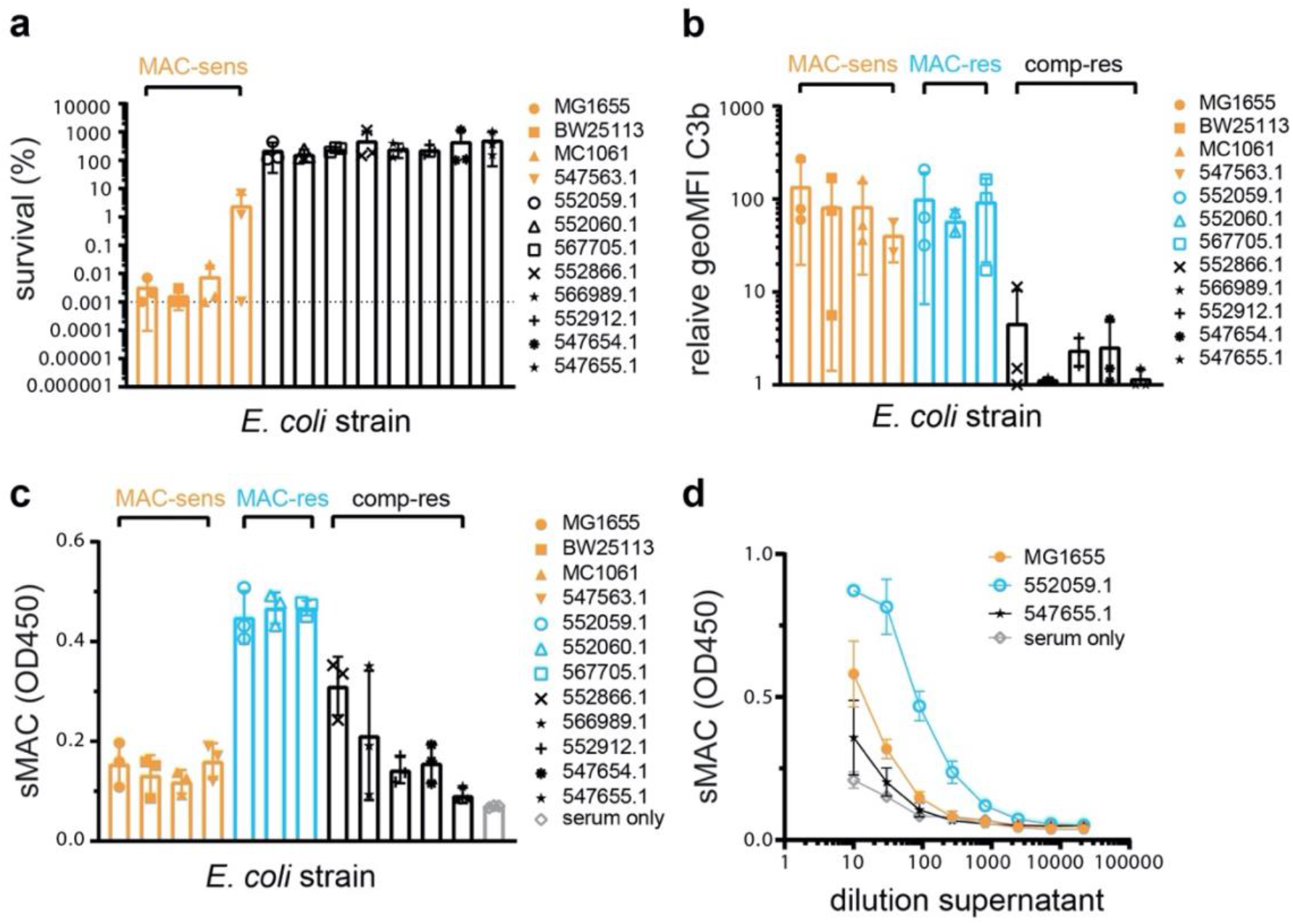
sMAC is primarily formed by MAC-resistant Gram-negative bacteria. *E. coli* strains (5×10^7^ bacteria/ml) were incubated in 5% pooled human serum. Bacterial viability was determined after 60 minutes by counting colony forming units (CFU’s) and calculating the survival compared to t=0. The horizontal dotted line represents the detection limit of the assay. b) *E. coli* strains (5×10^7^ bacteria/ml) were incubated in 10% C5-depleted serum. Bacteria were stained with AF488-labelled mouse monoclonal anti-C3b after 30 minutes and staining was measured by flow cytometry. The relative binding was calculated by normalizing the geoMFI to the geoMFI of unlabelled bacteria. c) sMAC was detected in the reaction supernatant (100-fold diluted) by ELISA after *E. coli* strains (5×10^8^ bacteria/ml) were incubated in 5% C6-depleted serum supplemented with C6-biotin for 60 minutes. Serum without bacteria (serum only) was taken as background control. d) sMAC was detected by ELISA for a dilution range of reaction supernatant collected in c for *E. coli* strain MG1655, 552059.1 and 547655.1. Orange strains are MAC-sensitive (MAC-sens), blue strains MAC-resistant (MAC-res) and black strains complement-resistant (comp-res). Flow cytometry data (b) are represented by individual geoMFI values of the bacterial population. Data represent individual values of three independent experiments with mean +/- SD.

We next wanted to see which *E. coli* strains formed sMAC in serum using an in-house sandwich ELISA. In short, sMAC in serum was captured using an antibody recognizing a neo-epitope of C9 when it is part of sMAC [21]. Next, C6 was detected with streptavidin-HRP, since we used C6-depleted serum supplemented with biotinylated C6. The specificity of our sMAC ELISA was validated using cobra venom factor (CVF) and C5 inhibitor eculizumab (**S2a**), which can form fluid-phase C5 convertases in serum [22]. All tested MAC-resistant strains efficiently formed sMAC in serum, whereas MAC-sensitive strains and most complement-resistant strains only generated slightly more sMAC than present in serum alone (**Fig. 1c**). Titration of the supernatant of MAC-resistant 552059.1 suggested that there was at least 5-fold more sMAC compared to MAC-sensitive MG1655 or complement-resistant 547654.1 (**Fig. 1d**). sMAC was formed in a complement-dependent manner, since C5-inhibitor OmCI and eculizumab prevented formation of sMAC (**S2b**). Finally, two MAC-resistant *Klebsiella* strains also formed more sMAC in serum compared to MAC-sensitive *Klebsiella* strains (**S2c**), suggesting that these findings can be translated to other Gram-negative species. Altogether, our data suggest that sMAC is primarily formed in serum by MAC-resistant Gram-negative bacteria.

### MAC-resistant *E. coli* strains more potently convert C5 in serum

Because conversion of C5 into C5b is crucial to initiate the assembly of sMAC, we wanted to know if conversion of C5 was also higher on all MAC-resistant *E. coli* strains compared to MAC-sensitive strains in serum. Western blotting of the serum supernatant confirmed that MAC-resistant strains converted all C5 in serum (**Fig. 2a**), whereas leftover C5 was still visible for MAC-sensitive strains. A sandwich ELISA was used to quantify the released C5a into serum supernatant, which revealed that MAC-resistant strains generated 5-fold more C5a compared to MAC-sensitive strains (**Fig. 2b**), which was comparable to the difference in sMAC (**Fig. 1d**). One MAC-sensitive strain (547563.1) also consumed more C5 and released more C5b into the supernatant compared to the other MAC-sensitive strains (**Fig. 2a, b**), although this difference was not detected by sMAC ELISA (**Fig. 1c**). Nonetheless, these data suggest that MAC-resistant *E. coli* strains also convert more C5 in serum.

**Figure 2.**
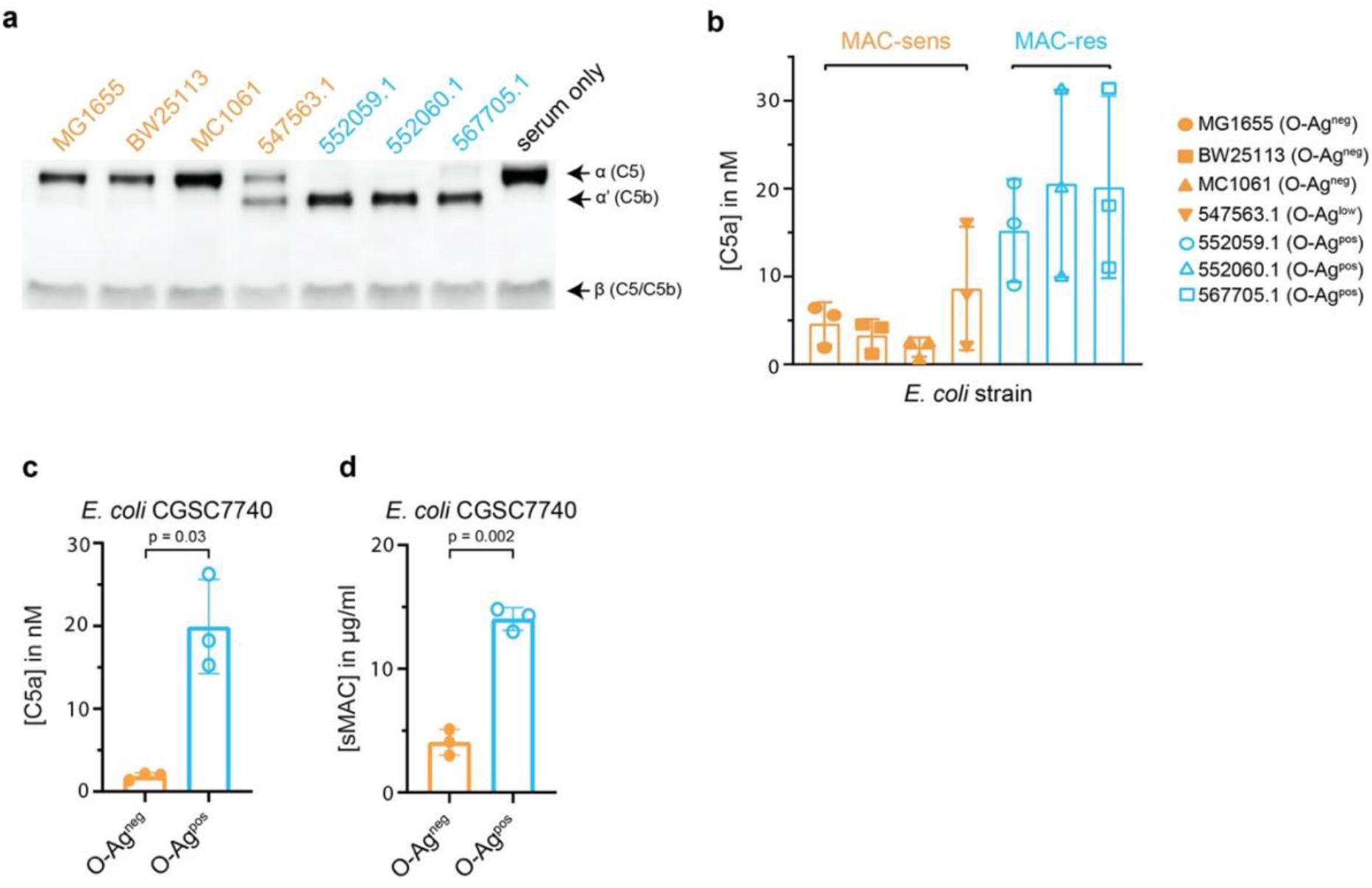
MAC-resistant *E. coli* strains convert more C5 in serum. *E. coli* strains (5×10^8^ bacteria/ml) were incubated in 5% pooled human serum and supernatant was collected by spinning down bacteria after 60 minutes. a) Representative Western blot for C5 of the supernatant. The upper band represents the α-chain of C5, the middle band of C5b (α’) and the lower band the β-chain of both C5 and C5b. Serum without bacteria (ser) was taken as control for the absence of C5 conversion. The Western blot is a representative of at least three independent experiments. b) C5a in the supernatant was quantified by ELISA. Orange strains are MAC-sensitive (MAC-sens) and blue strains are MAC-resistant (MAC-res). *E. coli* CGSC7740 wildtype without LPS O-Ag (O-Ag^neg^) and *wbbL*+ with LPS O-Ag (O-Ag^pos^) were also incubated in 5% pooled human serum to collect supernatant. C5a (c) and sMAC (d) in the supernatant were quantified by ELISA. ELISA Data represent individual values with mean +/- SD of three independent experiments. Statistical analysis was done using a paired two-tailed t-test (c and d) and relevant p-values are indicated in the figure.

### Expression of LPS O-Ag on *E. coli* increases C5a and sMAC generation in serum

We wondered if expression of lipopolysaccharide (LPS) O-Antigen (O-Ag) could explain this difference in C5 conversion. LPS O-Ag is an important constituent of the outer membrane of Gram-negative bacteria that has frequently been associated with MAC-resistance [23]. We have previously shown that the three tested MAC-resistant strains express O-Ag [20], whereas only one out of four MAC-sensitive strains (547563.1) does as well (shown in **S3a**, summarized in **Table 1**). Silver staining of O-Ag for *Klebsiella* strains suggested a comparable trend, showing little detectable O-Ag for MAC-sensitive strains compared to MAC-resistant strains (**S3b**). To more directly study if LPS O-Ag affects C5a generation and sMAC release, a MAC-sensitive *E. coli* K12 strain without O-Ag (O-Ag^neg^) was incubated in 5% human serum and compared with an isogenic MAC-resistant strain in which O-Ag expression is restored (O-Ag^pos^) [20]. Expression of O-Ag increased C5a generation in the supernatant 10-fold (**Fig. 2c**) and sMAC release 3.5-fold (**Fig. 2d**). C3b deposition on the bacterial surface was comparable both in the presence and absence of O-Ag in serum (**S3c**), suggesting that initial complement activation was not affected by the expression of O-Ag, similar to other MAC-resistant *E. coli* (**Fig. 1b**). Therefore, these data indicate that expression of O-Ag on *E. coli* can increase C5a generation and sMAC release in serum.

**Table 1.** Bacterial strains used in this study

| Strain | Origin | Amount of detectable LPS O-Ag |
| --- | --- | --- |
| *Escherichia coli* MG1655 | laboratory strain | absent [20] (also in **S3a**) |
| *Escherichia coli* BW25113 | laboratory strain | absent [20] (also in **S3a**) |
| *Escherichia coli* MC1061 | laboratory strain | absent [20] (also in **S3a**) |
| *Escherichia coli* 547563.1 | clinical isolate* | low# [20] (also in **S3a**) |
| *Escherichia coli* 552059.1 | clinical isolate* | high [20] (also in **S3a**) |
| *Escherichia coli* 552060.1 | clinical isolate* | high [20] (also in **S3a**) |
| *Escherichia coli* 567705.1 | clinical isolate* | high [20] (also in **S3a**) |
| *Escherichia coli* 566989.1 | clinical isolate* | high (**S3a**) |
| *Escherichia coli* 552912.1 | clinical isolate* | no (**S3a**) |
| *Escherichia coli* 552866.1 | clinical isolate* | high (**S3a**) |
| *Escherichia coli* 547654.1 | clinical isolate* | high (**S3a**) |
| *Escherichia coli* 547655.1 | clinical isolate* | high (**S3a**) |
| *Klebsiella variicola* 402 | clinical isolate* | low# (**S3b**) |
| *Klebsiella pneumoniae* 567880.1 | clinical isolate* | low# (**S3b**) |
| *Klebsiella pneumoniae* 567702.1 | clinical isolate* | high (**S3b**) |
| *Klebsiella pneumoniae* 567709.1 | clinical isolate* | high (**S3b**) |
| *Escherichia coli* CGSC7740 | laboratory strain | no [20] |
| *Escherichia coli* CGSC7740 *wbbL*+ | laboratory strain | high [20] |
| *Staphylococcus aureus* SH1000 | laboratory strain | n.a. |
| *Staphylococcus aureus* Wood46 | laboratory strain | n.a. |
| *Staphylococcus aureus* Newman | laboratory strain | n.a. |
| *Staphylococcus epidermidis* KV103 | clinical isolate* | n.a. |
| *Streptococcus agalactiae* COH-1 | clinical isolate* | n.a. |

\*All clinical isolates were obtained from the clinical Medical Microbiology department at the University Medical Center Utrecht. # detection of O-Ag was limited, but not absent. n.a. means O-Ag expression is not applicable, because these are Gram-positive bacteria that inherently do not express LPS.

### Binding of C8 and C9 triggers release of MAC precursors from MAC-resistant *E. coli*

We next studied at what stage of MAC assembly the nascent MAC is released from the bacterial surface. MAC pores assemble in a stepwise manner. C5b binds to C6 to form a stable C5b6 complex, which next binds C7 to anchor the C5b-7 complex to the membrane of bacteria [24]. Finally, binding of C8 inserts the nascent MAC into the bacterial cell envelope, and is more tightly inserted when C9 binds and polymerizes a transmembrane ring [25]. Release of C5b into the supernatant was therefore measured in the presence or absence of downstream MAC components for both MAC-resistant *E. coli* 552059.1 and MAC-sensitive *E. coli* MG1655. Bacteria were labelled with convertases in C5-depleted serum and washed as done previously [26]. Next, C5 and C6 were added in the presence or absence of downstream MAC components (**Fig. 3a**). Western blotting of the supernatant revealed that more C5 was converted into C5b for MAC-resistant 552059.1 compared to MAC-sensitive MG1655 (**Fig. 3b**), in line with **Fig 2b**. Western blotting also revealed that binding of C7 to the nascent MAC prevented release of C5b6 from the bacterial surface (**Fig. 3b**), as was already observed previously for MAC-sensitive MG1655 [27]. Binding of C7 also appeared to increase C5 conversion on MAC-resistant 552059.1 (**Fig. 3b**). However, binding of C8 to the nascent MAC triggered release of C5b from MAC-resistant 552059.1, even in the presence of final MAC component C9 (**Fig. 3b**). Binding of C8 also triggered some release of C5b from the surface of MAC-sensitive MG1655, but this was prevented when C9 was also present (**Fig. 3b**). Quantification of C5b6 by ELISA revealed that binding of C8 and C9 released 4-fold more C5b6 from the surface of MAC-resistant *E. coli* (**Fig. 3d**). These data suggest that binding of C8 and C9 to C5b-7 triggers release of the nascent MAC from MAC-resistant *E. coli*.

**Figure 3.**
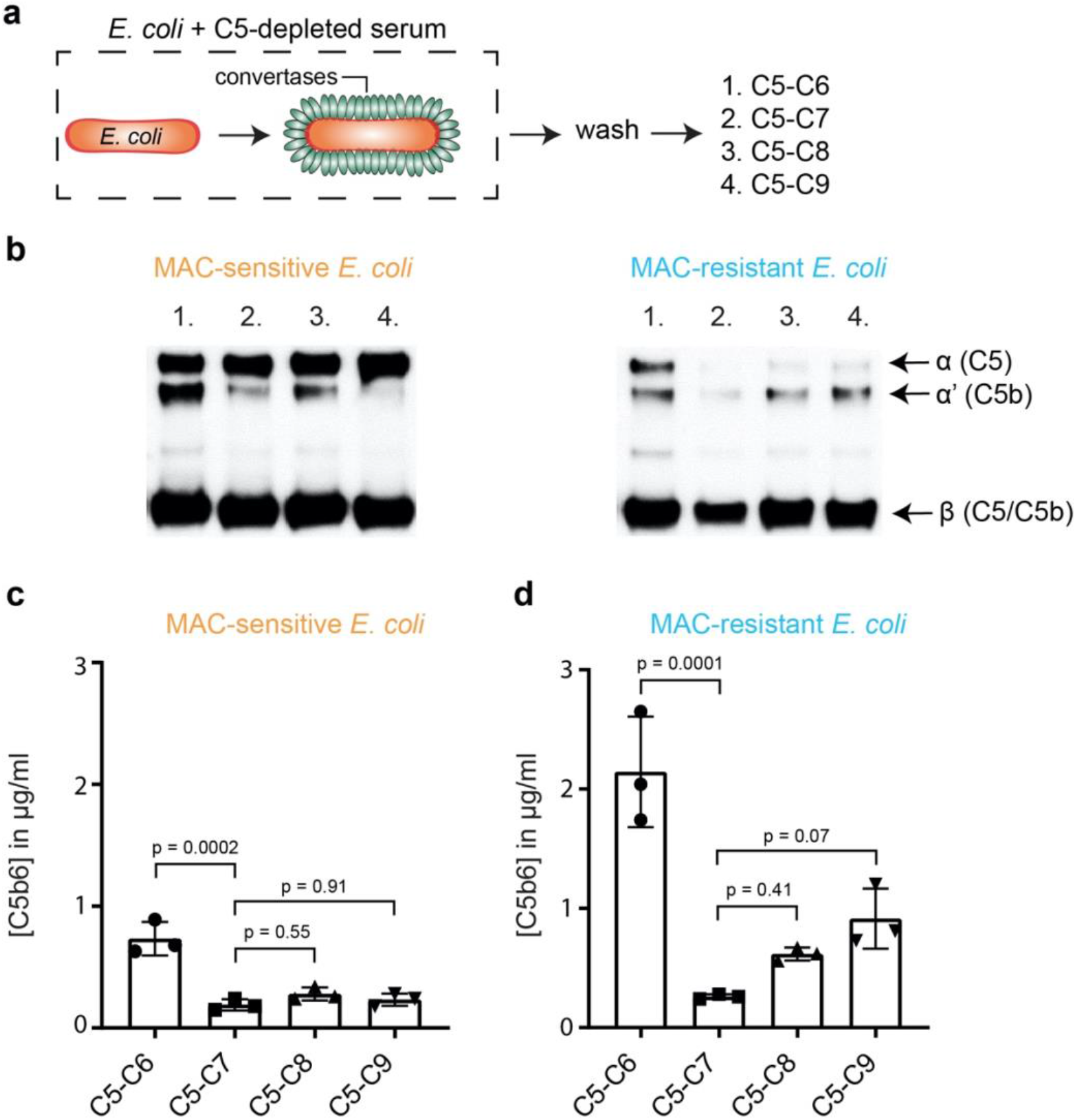
Binding of C8 and C9 release of MAC precursors from MAC-resistant *E. coli*. Schematic overview how *E. coli* (orange rods) MG1655 (MAC-senstive) and 552059.1 (MAC-resistant) were labelled with convertases (green ovals) in 10% C5-depleted serum. Next, bacteria were washed and bacteria (5×10^8^ bacteria/ml for b, 1×10^8^ bacteria/ml for c and d) were incubated with alternative pathway (AP) convertase components (5 µg/ml FB and 0.5 µg/ml FD) and 100 nM C5 and C6 (1); 100 nM C5, C6 and C7 (2); 100 nM C5, C6, C7 and C8 (3) or 100 nM C5, C6, C7, C8 and 1,000 nM C9 (4). The supernatant was collected after 60 minutes by spinning down bacteria. b) Western blot for C5 of the supernatant. The Western blot is a representative of at least three independent experiments. The upper band represents the α-chain of C5, the middle band of C5b (α’) and the lower band the β-chain of both C5 and C5b. C5b6 in the supernatant of MAC-sensitive MG1655 (c) and MAC-resistant 552059.1 (d) was quantified by ELISA. Dotted line represents the background OD450. ELISA data represent individual values of three independent experiments with mean +/- SD. Statistical analysis was done using an ordinary one-way ANOVA with Tukey’s multiple comparisons test (c and d) and relevant p-values are indicated in the figure.

### Release of MAC precursors from *E. coli* triggers lysis of bystander human erythrocytes, but is prevented by serum regulators vitronectin and clusterin

Although we measured the release of C5b (**Fig. 2**), the composition of the released complexes and their capacity to lyse cells remained unclear. Previous reports showed that complement activation on erythrocytes can cause MAC-dependent lysis of bystander cells that are not recognized by the complement system, so-called bystander lysis [28–30]. We next tested whether complement activation and subsequent release of MAC precursors from MAC-resistant *E. coli* can also cause bystander lysis. Therefore, *E. coli* were labelled with convertases in C5-depleted serum as done in **Fig. 2**. Next, these convertase-labelled bacteria were incubated with purified MAC components and unlabelled human erythrocytes to measure bystander lysis (**Fig. 4a**). This resulted in lysis of bystander erythrocytes for all MAC-resistant *E. coli* strains and MAC-sensitive 547563.1 (**Fig. 4b**), corresponding with the production of sMAC (**S4**). Lysis was prevented in the presence of C5 conversion inhibitor OmCI (**Fig. 4b**), suggesting that lysis was MAC-dependent. These data show that release of MAC precursors from *E. coli* can result in bystander lysis of human cells. However, when we studied bystander lysis in a human serum environment, we observed that the 552059.1 did not trigger bystander lysis of erythrocytes (**Fig. 4c**). Serum regulators Vn and Clu are known to scavenge and inactivate sMAC [31,32]. Indeed, both Vn and Clu inhibited bystander lysis of erythrocytes when MAC assembled on convertase-labelled *E. coli* 552059.1 (as described in **Fig. 4a**) at concentrations representative for 10% serum (**Fig. 4c**). SDS-PAGE revealed that Clu, but not Vn, prevents the formation of polymeric-C9 in the supernatant (**S5a**). This suggests that Vn and Clu interfere at different stages in the assembly of sMAC. Vn and Clu both specifically prevent lysis of bystander cells by MAC, since Vn and Clu did not inhibit binding of C9 (**S5b**), or MAC-dependent killing (**S5c**) when MAC was assembled by local conversion of C5 on convertase-labelled MAC-sensitive MG1655. Altogether, our data suggest that release of MAC precursors from *E. coli* can trigger bystander lysis of human erythrocytes, but that serum regulators Vn and Clu can inhibit this bystander lysis.

**Figure 4.**
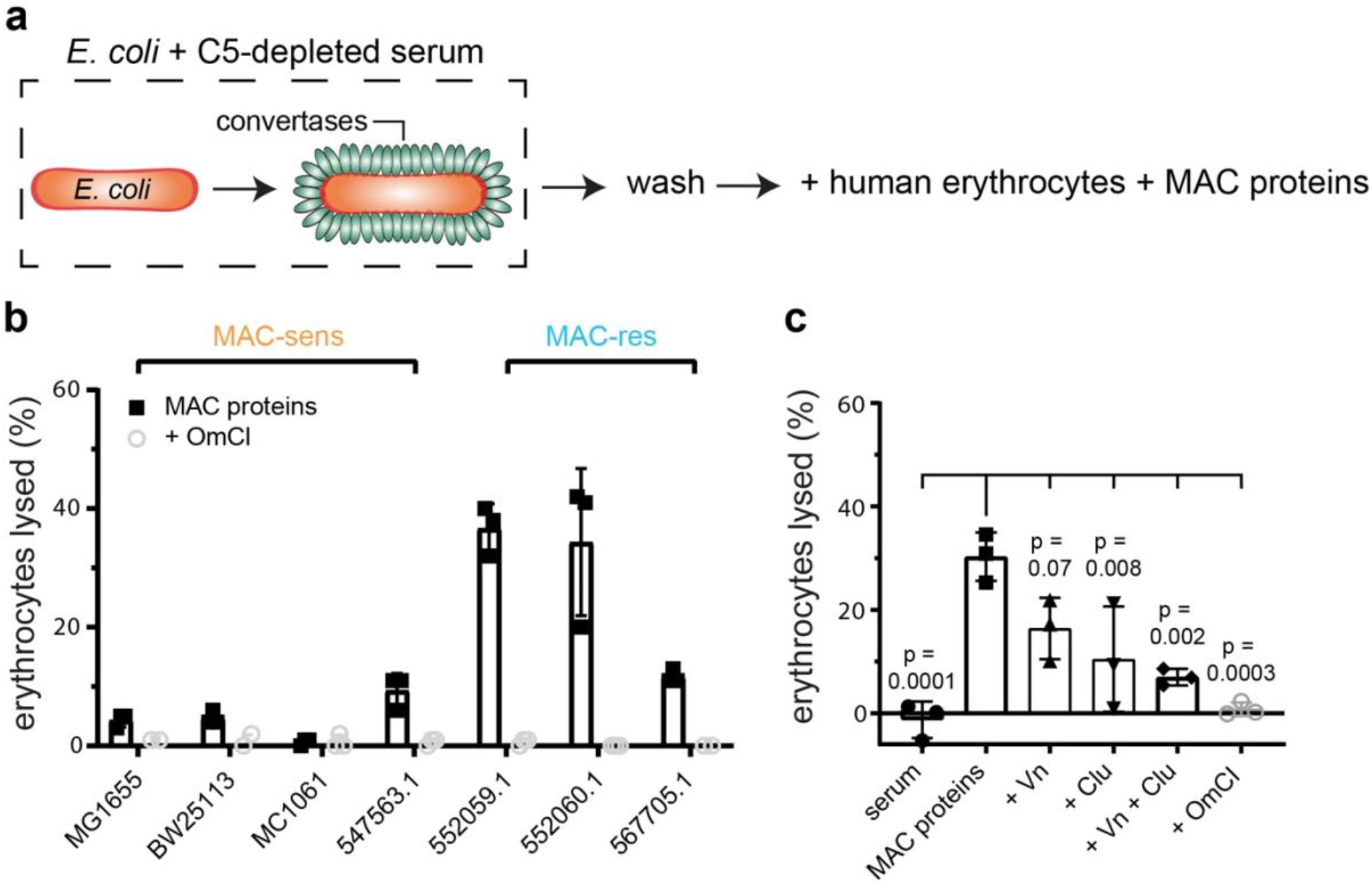
Release of MAC precursors from *E. coli* triggers lysis of bystander human erythrocytes, but is prevented by serum regulators vitronectin and clusterin. Schematic overview of the bystander lysis assay. *E. coli* strains (orange rods) were labelled with convertases (green ovals) in 10% C5-depleted serum and washed. Next, convertase-labelled bacteria (3.3×10^8^ per ml) were incubated with: human erythrocytes (1×10^8^ per ml), alternative pathway (AP) convertase components (5 nM FB and 20 nM FD) and MAC proteins (100 nM C5, 100 nM C6, 100 nM C7, 100 nM C8 and 500 nM C9). The supernatant was collected after 60 minutes by spinning down bacteria and erythrocytes and analyzed for the presence of haemoglobulin. The percentage of lysed erythrocytes was calculated by setting a buffer only control at 0% lysis and MilliQ control at 100% lysis. b) Bystander erythrocyte lysis for MAC-sensitive (MAC-sens) and MAC-resistant (MAC-res) *E. coli* strains. c) Bystander erythrocyte lysis for convertase-labelled MAC-resistant *E. coli* 552059.1 incubated with 10% pooled human serum, MAC proteins (30 nM C5, 30 nM C6, 30 nM C7, 30 nM C8 and 300 nM C9) or MAC components with 133 nM vitronectin (Vn), 133 nM clusterin (Clu) or 20 µg/ml C5 conversion inhibitor OmCI. Data represent individual values with mean +/- SD of three independent experiments. Statistical analysis was done using an ordinary one-way ANOVA with Tukey’s multiple comparisons test (c) and relevant p-values are indicated in the figure (all conditions compared with MAC proteins only).

### Gram-positive bacteria also release sMAC in serum

So far, our data indicate that on Gram-negative bacteria, sMAC is mainly released from MAC-resistant bacteria. Since sMAC is also increased in patient plasma that suffer from bacterial infections with Gram-positive bacteria [15,16], we also wanted to measure release of sMAC was released in serum by Gram-positive bacteria. Three *Staphylococcus aureus* strains (SH1000, Wood46 and Newman), one *Staphylococcus epidermidis* strain (KV103) and one *Streptococcus agalactiae* strain (COH-1) were incubated in serum. sMAC was formed in all five strains and was 3- to 5-fold higher compared to MAC-sensitive *E. coli* MG1655 (**Fig. 5a**), corresponding with MAC-resistant *E. coli* (**Fig. 1d**). Release of sMAC also coincided with generation of more C5a (**Fig. 5b**), although the increase compared to MAC-sensitive MG1655 was smaller for Gram-positive bacteria than for MAC-resistant *E. coli* (**Fig. 3a**). Altogether, these data indicate that Gram-positive bacteria also form sMAC in serum.

**Figure 5.**
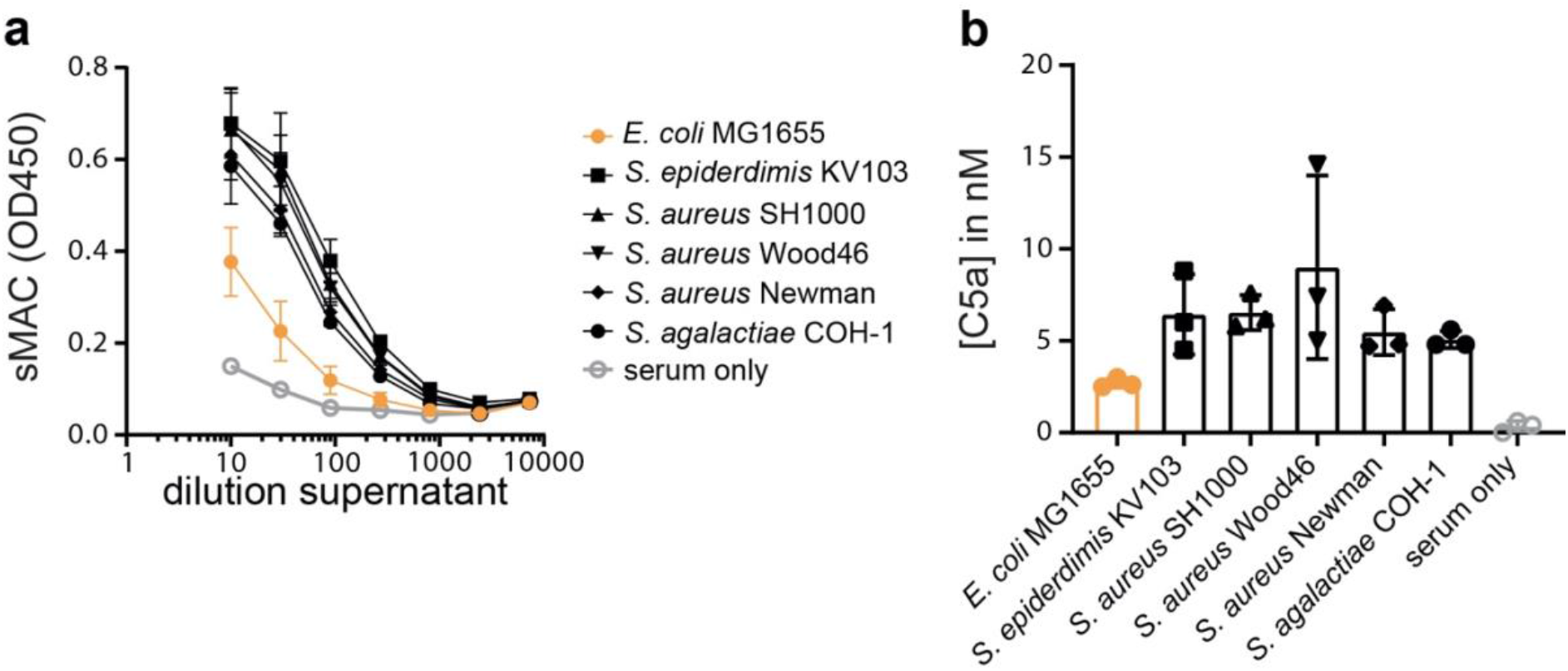
Gram-positive bacteria also release sMAC in serum. MAC-sensitive *E. coli* MG1655 and Gram-positive strains (5×10^8^ bacteria/ml) were incubated in 5% C6-depleted serum supplemented with C6-biotin. The supernatant was collected after 60 minutes by spinning down bacteria. Serum without bacteria (serum only) was taken as background control. a) sMAC was detected in a dilution range of reaction supernatant by ELISA. b) C5a in the supernatant was quantified by ELISA. Data represent individual values of three independent experiments with mean +/- SD.

### sMAC that is released from bacteria is a heterogeneous protein complex with different stoichiometries

Next, we aimed to define the molecular composition of sMAC generated when complement is activated on bacteria. sMAC was generated by incubating MAC-resistant *E. coli* 552059.1 in C6-depleted serum with His-tagged C6 and captured and isolated with HisTrap beads (**S6**). SEC was used to separate sMAC from monomeric-C6. The SEC profile of serum incubated with MAC-resistant *E. coli* shifted to much shorter elution times compared to non-activated serum (**Fig. 6a**, fractions B5-B12), indicating a mass shift. As a control, we analyzed commercially available sMAC, which is generated with zymosan particles in serum. Commercial sMAC eluted from the SEC column in the same fractions (B5-B12), indicating that these bacterial-eluate fractions contain sMAC. Blue-native PAGE (BN-PAGE) and subsequent Western blotting for C6 and C9 confirmed that these fractions contain sMAC (**Fig. 6b**). Compared to commercial sMAC, bacterial sMAC eluted somewhat later from the SEC column (**Fig. 6b**, most apparent in fractions B11 and B12). In addition, sMAC complexes generated by *E. coli* seemed to run further into the gel. These data suggest that bacterial sMAC complexes are have a different composition compared to commercial sMAC.

**Figure 6.**
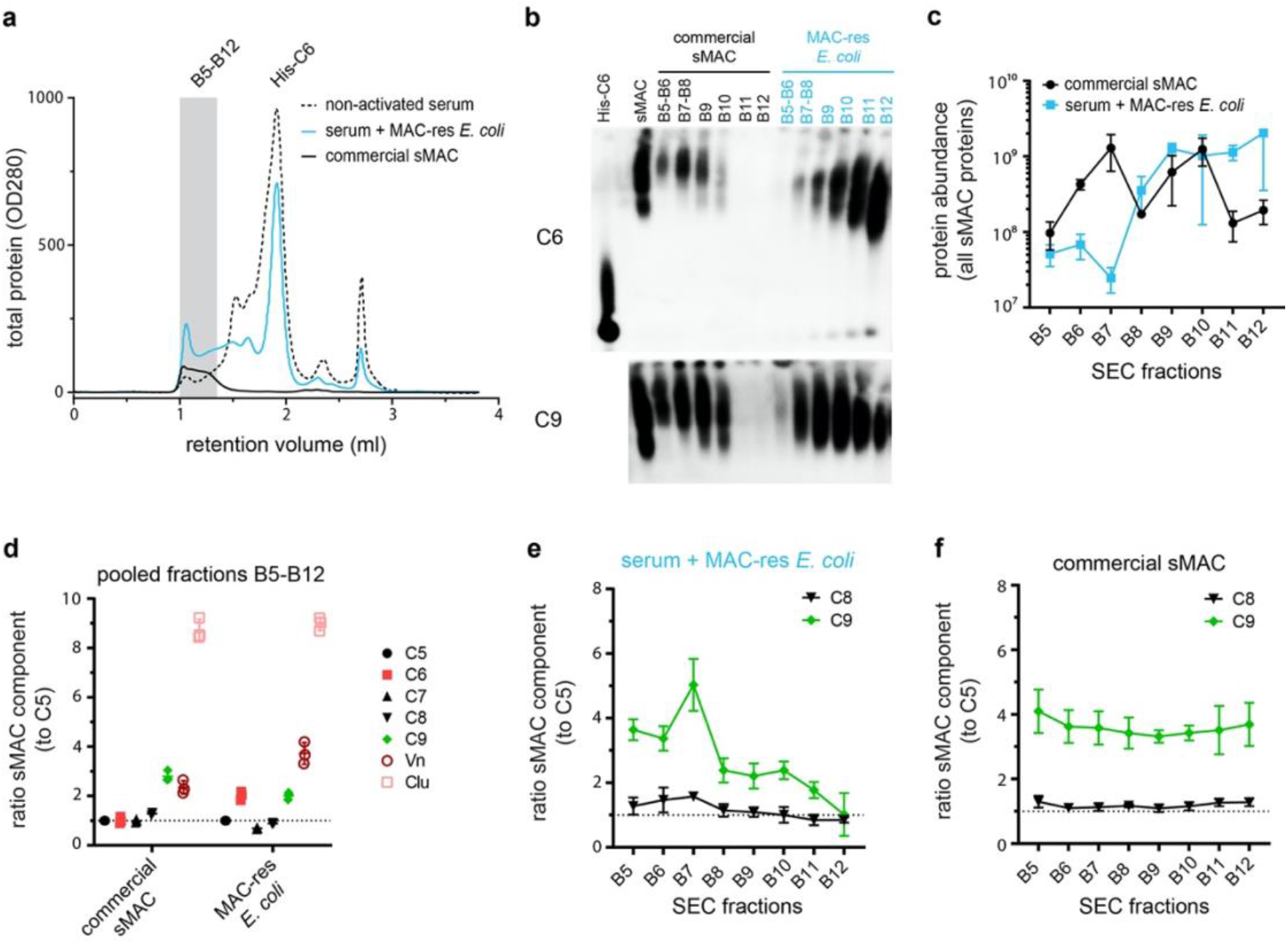
Isolation of sMAC using His-tagged C6 and analysis by BN-PAGE and LC-MS/MS. sMAC was generated by incubating MAC-resistant (MAC-res) *E. coli* 552059.1 in C6-depleted serum supplemented with His-tagged C6 (His-C6). sMAC in the supernatant was captured with HisTrap beads and eluted. a) Concentrated eluate was separated by SEC on a Superose 6 column and OD280 was measured to determine protein content (OD280). Eluate from serum without bacteria (non-activated serum) and 50 µg commercially available sMAC (Complement Technology) were analyzed as controls. b) BN-PAGE was performed with pooled (B5+B6 and B7+B8) or individual (B9-B12) SEC fractions and analyzed by Western blotting for C6 (above) and C9 (below). Black fractions represent SEC fractions commercially available sMAC, blue fractions represent SEC fractions of serum incubated with MAC-res *E. coli*. 2 µg of purified sMAC and His-C6 were loaded as control. c) The protein abundance of all sMAC components (iBAQ value) was determined by LC-MS/MS for individual SEC fractions (B5-B12). d) The ratio of all individual sMAC components to C5 was determined in a pooled sample of fraction B5-B12. e) In addition, the ratio of C8 and C9 to C5 was determined for each individual fraction separately for serum incubated with MAC-res *E. coli* and commercially available sMAC (f). The dotted line (d, e and f) represents a ratio of 1. The SEC profile and Western blot are representative of three independent experiments. LC-MS/MS data represent three individual digests of the same fraction with mean +/- SD that are representative for two independent experiments.

Menny *et al*. recently described that the commercially available sMAC used in our study consists of C5b-8, 1-3 copies of C9 and several copies of Vn and Clu, using proteomics, cross-linking mass spectrometry (MS) and cryo-electron microscopy (cryo-EM) [33]. Here, we compared the average composition of sMAC generated by MAC-resistant *E. coli* with commercial sMAC by profiling sMAC components with liquid chromatography-tandem mass spectrometry (LC-MS/MS). The total amount of sMAC components that were detected with LC-MS/MS in individual fractions (**Fig. 6c** and **S6b, c**) corresponded with BN-PAGE (**Fig. 6b**), suggesting that most bacterial sMAC eluted later from the SEC column than commercial sMAC. For both bacterial and commercial sMAC, sMAC components C5, C7, and C8 were present in equal amounts in a pooled sample of fractions B5-B12 (**Fig. 6d**). However, C6 and Vn were both 2-fold more abundant for bacterial sMAC (**Fig. 6d**). The increased ratio of C6 is likely explained by the fact that not all monomeric-C6 could be separated during SEC (as was visible in fraction B9-B12 by BN-PAGE in **Fig. 6b**). Surprisingly, bacterial sMAC contained on average less C9 per C5 (**Fig. 6d**, ratio of 2:1) than commercially available sMAC (**Fig. 6d**, ratio of 3:1). The relative amount of C9 per C5 (**Fig. 6e**), but not other sMAC components (**S6d**), decreased in fractions that eluted later from the SEC column for bacterial sMAC. This was not observed for commercially available sMAC (**Fig. 6f** and **S6e**). These data suggest sMAC generated by bacteria contains more complexes with less C9 compared to commercial sMAC.

Finally, we assessed the relative abundance of sMAC components for sMAC that was generated by Gram-positive bacteria. *S. aureus* Wood46 was incubated with human serum and the supernatant was separated by SEC (without isolating sMAC via HisTrap beads), using non-activated serum as control. By profiling sMAC components in individual fractions by LC-MS/MS, we found that in non-activated serum these components elute later in the SEC profile, corresponding to monomeric or low molecular weight complexes (**Fig. 7**). Upon activation, the profiles of all sMAC components largely co-elute and are shifted to fractions that correspond to the higher mass range. In fact, close to no monomeric MAC components were detected after incubation with bacteria, suggesting that all available MAC components were incorporated into hetero-oligomeric sMAC complexes. We did not observe any sMAC without Vn and Clu in the bacterial activated sample, since these would be located around the 664 kDa molecular weight marker, suggesting that all sMAC complexes were bound to multiple Vn and Clu molecules (corresponding with the relative abundance in **Fig. 6d**). The complexes correlate with a ratio of 1-3 C9 molecules and an average of 2 C9 molecules to C5b-8, with Clu as the most abundant regulator bound to sMAC. Altogether, these data confirm that sMAC released from bacteria is a heterogeneous assembly comprising a single copy of C5b-8 together with multiple copies of C9, Clu and Vn in a mixture of stoichiometries.

**Figure 7.**
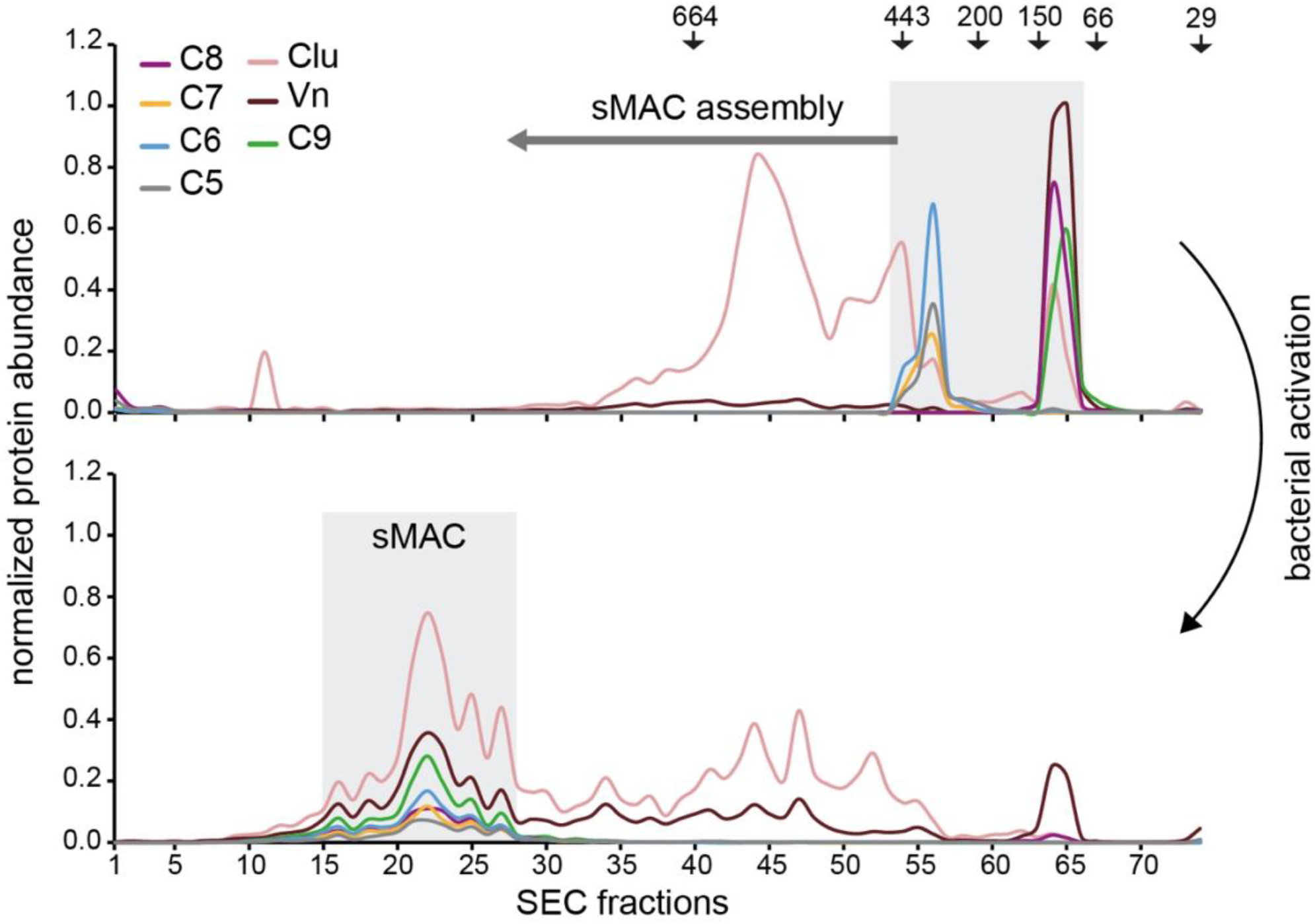
sMAC that is released from bacteria is a heterogeneous protein complex with different stoichiometries. MS profiling of sMAC components in serum supernatant incubated for 3 hours at 37 °C with (bottom, activated) and without (top, nonactivated) *S. aureus* Wood46. Serum was separated by SEC and the protein abundance (normalized iBAQ values) in each fraction was determined by LC-MS/MS. The grey boxes indicate the elution of sMAC components in nonactivated and activated serum. The arrows on the top indicates elution of molecular weight (kDa) markers.

## Discussion

Although plasma levels of sMAC are frequently increased during bacterial infections [15,16], it remains largely unknown how sMAC is formed during infections and what the complex represents. Here, we show that sMAC is an inactivated complex that is released from bacteria during complement activation. We show that sMAC is primarily released from MAC-resistant bacteria, including Gram-positive bacteria. Surprisingly, these MAC-resistant bacteria also potently activate the complement cascade and convert more C5 than MAC-sensitive bacteria.

These findings suggest that detection of sMAC in human serum indicates potent C5 conversion by MAC-resistant bacteria. Increased C5 conversion has previously been associated with MAC-resistance on Gram-negative bacteria [34,35], but this has not yet been directly linked to the detection of sMAC in human serum. Moreover, our study extends on these insights by showing that potent C5 conversion by MAC-resistant *E. coli* is linked to the expression LPS O-Ag. How expression of LPS O-Ag results in more potent conversion of C5 compared to MAC-sensitive strains remains unclear. Grossman *et al*. have previously demonstrated that linking *Salmonella* LPS O-Ag to sheep erythrocyte membranes can increase C3 consumption [36]. However, in our study, no difference was detected in C3 conversion and C3b deposition, suggesting that initial complement activation was comparable. For the conversion of C5, the density of C3b is known to be important [37,38]. Although C3b deposition was comparable between MAC-sensitive and MAC-resistant *E. coli*, we cannot exclude that local densities of C3b might differ, which could explain a difference in C5 conversion. In addition, the serine protease Bb that is bound to C3b and forms the convertases of the alternative pathway (AP) can be stabilized by properdin [39–42]. Differences in binding of properdin could cause a difference in the stability of AP convertases between strains. This is also supported by our previous study [26], in which we have shown that pre-labelling of bacteria in C5-depleted serum and subsequent washing, as done here, primarily results in functional AP convertase. Finally, the presence of downstream MAC components specifically enhanced C5 conversion for a MAC-resistant *E. coli* strains, suggesting that assembly of MAC could also affect further C5 conversion.

Our study also provides molecular insight into how sMAC is formed when complement is activated on bacteria. Although it was generally believed that sMAC is formed on the bacterial surface and then released [19], it was still unclear if the complete complex is formed on the bacterial surface. Our findings suggest that sMAC is initially formed on the bacterial surface, but released as the nascent MAC further assembles. Especially binding of C8 and C9 to C5b-7 triggered release of the nascent MAC from the bacterial surface. Our MS data show that most sMAC complexes ultimately contain on average 1 to 3 C9 molecules and a varying amount of Vn and Clu copies, which implies that sMAC further assembles in solution. Joiner *et al*. have previously observed that binding of C8 to C5b-7 on a MAC-resistant *Salmonella Minnesota* strain triggered release of C5b-8 [43]. Although binding of C8 to soluble C5b-7 prevents binding to membranes [44], our data suggest that released MAC precursors can still bind to bystander cells and form lytic MAC pores. It is possible that C5b-8 can immediately bind bystander cells in our assays, which prevents C5b-7 from entering a soluble state. However, we cannot exclude that trace amounts of intermediate MAC precursors, such as C5b6, that are also released from the bacterial surface and can still form enough MAC pores to lyse bystander erythrocytes.

Since C8 and C9 are the MAC components that insert into membranes [6,45], our findings suggest that the nascent MAC is released because it is less capable of inserting into the bacterial cell envelope of MAC-resistant bacteria. Why MAC insertion is impaired on MAC-resistant bacteria remains unclear. On Gram-positive bacteria, it is believed that the dense peptidoglycan layer prevents insertion of MAC into the cytoplasmic membrane [12]. On Gram-negative bacteria, MAC-resistance and improper insertion of MAC into the bacterial outer membrane have previously been linked to the expression and length of the LPS O-Ag [23]. We here show that the expression of LPS O-Ag directly increases the generation and release of sMAC. LPS O-Ag could prevent MAC proteins from inserting into hydrophobic patches of the bacterial outer membrane, ultimately resulting in release of sMAC. This is in line with our previous study, which demonstrated that expression of O-Ag impairs polymerization of C9 at the bacterial surface [20]. The presence of LPS O-Ag could also prevent binding of complement-activating antibodies that bind to epitopes close to the bacterial surface [46]. Instead, antibodies that recognize the O-Ag could activate complement further away from the surface, which prevents insertion of MAC proteins into OM and results in release of sMAC.

Findings in this study also highlight the importance of Vn and Clu in preventing bystander lysis of host cells when bacteria activate complement. In the absence of Vn and Clu, release of MAC precursors from bacteria resulted in lysis of bystander human erythrocytes. Importantly, Vn and Clu did not prevent MAC-dependent killing of the target bacterium, suggesting that Vn and Clu specifically inhibit bystander lysis. This bystander lysis has been studied using sensitized and unsensitized erythrocytes in the past [28–30], but relatively few reports have studied bystander lysis in a bacterial context [47,48]. In these reports, complement activation on *Streptococcus pneumoniae* [47] and *Moraxella catarrhalis* [48] caused bystander lysis of chicken erythrocytes. We here extend on these findings showing that human erythrocytes are also sensitive to bystander lysis.

This raises the question if bystander lysis is a clinically relevant process during bacterial infections. Under physiological conditions, Vn and Clu are present in a 2-to 8-fold molar excess compared to MAC components in serum, favouring inactivation of released MAC over binding to a bystander cell membrane [15]. Correspondingly, we do not observe bystander lysis in a serum of healthy donors in our study. However, Willems *et al*. have recently reported that Clu is decreased in plasma of children suffering from bacterial infections [49]. Vn and Clu polymorphisms have also been identified that impair their potency in scavenging and inactivating sMAC precursors [50,51], which could predispose to host cell damage by bystander lysis. For both polymorphisms, patients suffered from recurrent complement-mediated hemolytic uremic syndrome, and in case of the Vn polymorphism, this was even associated with recurrent *E. coli* infections. Moreover, both Gram-negative and Gram-positive bacterial pathogens are known to recruit Vn to their surface [52,53]. For Gram-negative bacteria, this is thought to prevent MAC-dependent killing [54,55], but for Gram-positive bacteria, the purpose of recruiting Vn and Clu has remained elusive. Altogether, our study therefore suggests a potential role of bystander lysis during bacterial infections that merits further investigation.

Finally, we demonstrate that sMAC generated by bacteria is a heterogeneous mixture of alike protein complexes with different stoichiometries. sMAC is believed to consist of C5b-7, C8, multiple copies of C9 and several copies of Vn and Clu [15,56]. This was recently supported by Menny *et al*. [33], who revealed that sMAC generated by zymosan particles in serum contains at least 1-3 copies of C9 [33]. Our data suggest that sMAC generated by bacteria is present in similar stoichiometries, but on average, contains less C9 molecules, especially for sMAC generated by MAC-resistant *E. coli*. This suggests that the stoichiometry of sMAC could depend on the target cell that activates complement. However, it is important to note the possibility that complexes with less C9 are lost during purification of commercially available sMAC. Our data also show the presence of multiple copies of Vn and Clu in sMAC generated by bacteria, with Clu being the most abundant chaperone. These findings suggest that Vn and Clu can efficiently capture all sMAC that is released in serum. This is in line with the recent structure of sMAC by Menny *et al*. [33]. That structure also revealed that Clu binds and traps the terminal C9 in an intermediate conformation, thereby preventing further C9 polymerization [33]. This is in line with our results showing that Clu, and not Vn, is able to inhibit C9 polymerization. Compared to commercial sMAC, more Vn was detected in sMAC generated by bacteria. It is unclear if this is caused by a difference in the target cell that activates complement, or a difference in the serum that was used to generate sMAC. The concentration of Vn in serum can vary largely between individuals [15] and could be responsible for the observed difference.

Altogether, we show that sMAC represents an inactivated complex that is primarily released from MAC-resistant bacteria that potently activate complement. These findings are clinically relevant as they provide insight into how sMAC can be used as a biomarker during bacterial infections. These insights could also help further understand the role of complement activation during bacterial infections in pathogenesis.

## Materials & methods

### Serum and complement proteins

Pooled human serum was obtained from healthy volunteers as previously described [57]. Serum depleted of complement components C5 or C6 was obtained from Complement Technology. Cobra venom factor (CVF) was obtained from Quidel. Preassembled C5b6, C8 and sMAC (SC5b-9) were obtained from Complement Technology. His-tagged C5, C6, C7 and factor B (FB) were expressed in HEK293E cells at U-Protein Express as described previously [27]. Factor D (FD) and OmCI were produced in HEK293E cells at U-Protein Express and purified as described before [58]. To produce fluorescently labelled C9, C9-3xGGGGS-LPeTG-6xHis was recombinantly expressed in Expi293F cells and site-specifically labelled with Cy5 via C-terminal sortagging as done in [26]. Biotinylated C6 was produced in a similar manner, by expressing and isolating C6-LPeTGG-6xHis (previously described in [27]) and subsequent C-terminal sortagging with GGGK-biotin (kindly provided by Louris Feitsma, Department of Crystal and Structural Chemistry, Bijvoet Insitute). Eculizumab was kindly provided by Genmab. Vitronectin (plasma isolated) was obtained from Advanced Biomatrix and recombinantly expressed human clusterin from R&D systems. Monoclonal mouse-anti C3b (bH6, kindly provided by Peter Garred) was randomly labelled with NHS-Alexa Fluor 555 (AF555, Thermofisher) as done previously [26]. The concentrations of MAC components in 100% serum are ∼ 375 nM C5, 550 nM C6, 600 nM C7, 350 nM C8 and 900 nM C9.

### Bacterial growth

Bacterial strains that were used in this study are shown in **Table 1**. CGSC7740 wildtype (O-Ag^neg^) and *wbbL+* (O-Ag^pos^) were kindly provided by Benjamin Sellner (Biozentrum, University of Basel). In the *wbbL*+ strain, an IS5-element that inactivates the *wbbL* gene is removed. This restores expression of an essential rhamnose transferase wbbL that is required for the expression of O-Ag [20,59]. CGSC7740 *wbbL*+ was constructed by replacing the IS5-element present in the *wbbL*+ gene with a *sacB-kan* cassette to select for kanamycin-resistance. The *sacB-kan* cassette was then replaced with wildtype *wbbL* without the IS5-element and selected by counter selection on sucrose.

Bacteria were plated from glycerol stocks on blood agar plates. Single colonies were picked and grown overnight at 37 °C in shaking conditions (600 rpm). Gram-negative bacteria were grown in lysogeny broth (LB). *S. aureus* were grown in Todd Hewitt Broth (THB) and *S. epidermidis* in Trypticase Soy Broth (TSB). *S. agalactiae* COH-1 was grown in THB in non-shaking conditions and at 5% CO2. The next day, subcultures were grown by diluting at least 1/30 and these were grown to mid-log phase (OD600 between 0.4 – 0.6). Once grown to mid-log phase, bacteria were washed by centrifugation three times (11,000 rcf for 2 minutes) and resuspended to OD 1.0 (1 × 10^9^ bacteria/ml, validated by flow cytometry for all individual bacterial strains) in RPMI (Gibco) + 0.05% human serum albumin (HSA, Sanquin).

### Complement activation and killing in serum

To activate complement on bacteria, bacteria (the amount is specified in figure legends) were incubated in 5% pooled human serum for 60 minutes at 37 °C. For sMAC ELISA’s with C6-biotin, C6-depleted serum was used supplemented with 28 nM of C6-biotin. For ELISA’s and Western blotting, supernatant was collected by spinning down bacteria at 11,000 rcf for 2 minutes. Blocking of C5 conversion in serum was done with 6 µg/ml OmCI with and without 6 µg/ml eculizumab.

### Convertase labelling and purified MAC formation

Bacteria were labelled with convertases in C5-depleted serum as validated previously [26]. In short, bacteria (5 × 10^8^ bacteria/ml) were incubated with 10% C5-depleted serum for 30 minutes at 37 °C, washed three times (11,000 rcf for 2 minutes) and resuspended in RPMI-HSA. Bacteria were counted by flow cytometry to ensure that bacterial concentrations were comparable between different strains after convertase-labelling. Convertase-labelled bacteria (concentrations specified in figure legends) were next incubated with MAC components (concentrations specified in figure legends) and AP convertase components (50 nM FB and 20 nM FD) for 60 minutes at 37 °C. To collect the supernatant for ELISA and Western blotting, bacteria were spun down at 11,000 rcf for 2 minutes.

### Bacterial viability

Bacterial viability was assessed by determining colony forming units (CFU’s). A serial dilution was made in PBS (100, 1,000, 10,000 and 100,000-fold) and plated in duplicate on LB agar plates. After overnight incubation at 37 °C, colonies were counted and the corresponding concentration of CFU/ml was calculated. Survival was calculated by dividing the CFU/ml in the sample by the CFU/ml at t=0.

### Flow cytometry

To measure killing by flow cytometry, 2.5 µM of Sytox Blue Dead Cell stain (Thermofisher) was added during the assay to measure IM damage. To measure C3b deposition by flow cytometry, bacteria (∼5 × 10^7^ bacteria/ml) were stained after labelling with convertases with 3 µg/ml AF555 labelled mouse-anti C3b for 30 minutes at 4 °C. Finally, bacterial samples were diluted to ∼1 × 10^6^ bacteria/ml in RPMI-HSA and subsequently analyzed in a MACSquant flow cytometer (Miltenyi) for forward scatter (FSC), side scatter (SSC), Sytox, AF555 and Cy5 intensity. Flow cytometry data was analyzed in FlowJo V.10. Bacteria were gated on forward scatter and side scatter.

### Complement activation product ELISA’s

Serum supernatants were diluted (as specified in figure legends) in PBS + 0.05% Tween (PBS-T) supplemented with 1% bovine serum albumin (BSA). Next, sample dilutions were analyzed for the presence of complement activation products C3a, C5a, sMAC and C5b6 via enzyme-linked immunosorbent assays (ELISA) on Nunc Maxisorp ELISA plates. For C5a, a sandwich-ELISA kit was used (R&D Systems, DY2037), which includes two mouse monoclonal C5a antibodies that specifically detect in C5a and not native C5.

For C3a, sMAC and C5b6, plates were coated overnight at 4 °C with 1 µg/ml of coating antibody. For C3a this was mouse monoclonal anti-C3a (Hycult), for C5b6 this was monoclonal mouse IgG1 anti human C6 (Quidel) as used previously [27] and for sMAC this was mouse monoclonal aE11 directed against a neo-epitope of C9 in sMAC (kindly provided by T. Mollness and P. Garred). Blocking was next performed with PBS-T + 4% BSA for 60 minutes at RT. Sample dilutions were next bound for 60 minutes at RT. Primary staining was performed for C3a with 1:2,000 rabbit anti-C3a (Calbiochem), for C5b6 with 1:500 dilution of goat-anti human C5 serum (Complement Technology) and for sMAC with 1 µg/ml biotinylated monoclonal anti-C7 (clone F10, described in [60] and kindly provided by Wioleta Zelek). Samples that were prepared in C6-depleted serum supplemented with C6-biotin were directly stained with 1:5,000 HRP-conjugated streptavidin to detect sMAC. Otherwise, secondary staining was performed for C3a with 1:5,000 HRP-conjugated polyclonal goat antisera against rabbit IgG (Southern Biotech), for C5b6 with a 1:5.000 of HRP-conjugated donkey antisera against goat IgG (H+L) (Southern Biotech) and for sMAC with 1:5,000 HRP-conjugated streptavidin (Southern Biotech). Finally, fresh tetramethylbenzidine (TMB) was added for development and the reaction was stopped with 4N sulfuric acid to measure OD450.

At each step for all ELISA’s, 50 µl was added per well, antibodies were diluted in PBS-T + 1% BSA and incubation was done for 60 minutes at RT (except for coating). In between steps, wells were washed three times with PBS-T in between each step. Quantification of C3a, C5a, C5b6 and sMAC were done by interpolation with a standard curve of purified C3a-desarginine, C5a (Bachem), C5b6 and sMAC/SC5b-9 (Complement Technology).

### C5b Western blots

Bacterial supernatants were collected as described above and the cell pellets were also collected. Samples were diluted 1:1 in 2x reducing SDS sample buffer (0.1M Tris (pH 6.8), 39% glycerol, 0.6% SDS and bromophenol blue) supplemented with 50 mg/ml dithiothreitol (DTT) and incubated at 95 °C for 5 minutes. Samples were run on a 4-12% Bis-Tris gradient gel (Invitrogen) for 60 minutes at 200V. Proteins were next transferred with the Trans-Blot Turbo Transfer system (Bio-Rad) to 0.2 µM PVDF membranes (Bio-Rad). Initially, samples were blocked with PBS supplemented with 0.1% Tween-20 (PBS-T) and 4% dried skim milk (ELK, Campina) for 60 minutes at 37 °C. Primary staining was performed with a 1:500 dilution (∼ 80 µg/ml) of polyclonal goat-anti human C5 (Complement Technology) in PBS-T supplemented with 1% ELK for 60 minutes at 37 °C. Secondary staining was performed with a 1:10,000 dilution of HRP-conjugated pooled donkey antisera against goat IgG (H+L) (Southern Biotech) in PBS-T supplemented with 1% ELK for 60 minutes at 37 °C. In between all steps and after the final staining, membranes were washed three times with PBS-T. Finally, membranes were developed with Pierce ECL Western Blotting Substrate (Thermo Scientific) for 1 minute at RT and imaged on the LAS4000 Imagequant (GE Healthcare).

### Bystander lysis assay

Human erythrocytes were collected from heparin-sulfate treated human blood of healthy donors. Erythrocytes were washed at 1,000 rcf for 5 minutes for three times with PBS. The packed erythrocyte pellet was resuspended in Veronal buffered saline (2 mM Veronal, 145 mM NaCl, pH = 7.4) supplemented with 0.1% BSA and 2.5 mM MgCl2 (VBS+) and diluted to 4% (∼1×10^8^ erythrocytes/ml). Erythrocytes (1×10^8^ erythrocytes/ml) were next incubated with convertase-labelled bacteria (5×10^8^ bacteria/ml) and 100 nM C5-C8, 300 nM C9, 50 nM FB and 20 nM FD (unless specified otherwise in figure legends) for 60 minutes at 37 °C. Supernatant of the reaction was collected by spinning down the plate (1,250 rcf for 5 minutes) and the supernatant was next diluted 1:3 in MilliQ (MQ). Haemoglobulin release was measured by measuring the absorbance at OD405 nm. The percentage of lysed erythrocytes was calculated by setting a buffer only control at 0% lysis and a MQ control at 100% lysis. Blocking of C5 conversion in serum was done with 15 µg/ml OmCI.

### Polymeric-C9 detection by SDS-PAGE

Reaction supernatants were resuspended and diluted 1:1 in 2x SDS sample buffer supplemented with 50 mg/ml DTT and incubated at 95 °C for 5 minutes. Samples were run on a 4-12% Bis-Tris gradient gel (Invitrogen) for 75 minutes at 200V. Gels were imaged for 10 minutes with increments of 30 seconds on the LAS4000 Imagequant (GE Healthcare) for in-gel Cy5 fluorescence. Monomeric-C9 (mono-C9) and polymeric-C9 (poly-C9) were distinguished by size, since mono-C9 runs at 63 kDa and poly-C9 is retained in the comb of the gel.

### HisTrap isolation of sMAC with His-tagged C6

*E. coli* bacteria (5×10^8^ bacteria/ml) were incubated in 10% C6-depleted serum supplemented with 50 nM His-tagged C6 in a total volume of 1 ml for 60 minutes at 37 °C. Serum supernatant was collected as described above and incubated with a pellet of 900 μl Dynabeads™ His-Tag Isolation & Pulldown (Invitrogen) equilibrated in wash buffer (50 mM Tris, 300 mM NaCl, 10 mM imidazole, pH 7.8) on a tube rotator for 90 minutes at 4°C. Beads were separated using a magnet to collect the bead supernatant. Beads were washed three times in wash buffer and next His-tagged proteins were eluted with elution buffer (50 mM Tris, 300 mM NaCl, 300 mM imidazole, pH 7.8) on a tube rotator for 30 minutes at 4°C. Beads were separated using a magnet to collect the eluate. The eluate was finally filtered through a 0.22 µm filter and concentrated in a 100 kDa Amicon tube. sMAC was next separated from free leftover proteins by size exclusion chromatography (SEC) on a Superose 6 Increase column with PBS. 50 μl fractions were collected and used for subsequent analyses.

### BN-PAGE and Western blot

Samples were diluted 1:1 in 2x NativePAGE sample buffer (Invitrogen). SDS-PAGE samples were run on a 4–12% Bis-Tris gradient gel (Invitrogen) for 75 minutes at 200V. Samples were run on a NativePAGE 3-12% Bis-Tris gradient gel (Invitrogen) for 3 hours at 150V, after which the gel was destained overnight with demi water. Proteins were transferred to 0.2 µM PVDF membranes with the Trans-Blot Turbo Transfer system (Bio-Rad). Membranes were blocked in PBS/0.1% Tween (PBS-T) with 4% ELK (Campina) for 45 minutes at 37°C. Primary detection antibodies (polyclonal goat anti-human C9 from Complement Technology) were diluted 1:500 in PBS-T/1% ELK and incubated for 45 minutes at 37°C. Secondary detection antibody (HRP-conjugated donkey anti-goat IgG from Southern Biotech) was diluted 1:10,000 in PBS-T/1% ELK and incubated for 45 minutes at 4°C. In between each step, membranes were washed three times with PBS-T. The staining was developed with Pierce ECL Western Blotting Substrate (Thermo Scientific) for 1 minute at RT. Images were made using the LAS4000 Imagequant (GE Healthcare).

### SEC of activated and nonactivated serum samples

200 µl *S. aureus* Wood 46 (1.5×10^9^ bacteria/ml) in PBS was pelleted and resuspended in 250 µl serum for 3 h at 37°C while shaking. Bacteria were spun down at 11,000 rcf rpm for 3 min. The supernatant was collected and the centrifugation step repeated to spin down remaining bacteria. The supernatant was then kept on ice and filtered through a 0.22 µm filter. SEC separation of serum samples was done using an Agilent 1290 Infinity HPLC system (Agilent Technologies) consisting of a vacuum degasser, refrigerated autosampler with a 100 µl injector loop, binary pump, thermostated two-column compartment, auto collection fraction module, and multi-wavelength detector. The dual-column set-up, comprising a tandem Yarra™ 4000-Yarra™ 3000 (SEC-4000, 300 × 7.8 mm i.d., 3 µm, 500 Å; SEC-3000, 300 × 7.8 mm i.d., 3 µm, 290 Å) two-stage set-up. Both columns were purchased from Phenomenex. The columns were cooled to 17°C while the other bays were chilled to 4°C to minimize sample degradation. The mobile phase buffer consisted of 150 mM AMAC in water and filtered using a 0.22 µm disposable membrane cartridge (Millipore) before use. Approximately 1.25 mg of serum protein (activated and nonactivated fresh serum) was injected per run. The proteins were eluted using isocratic flow within 60 min, and the flow rate was set to 500 µl min-1. In total, 74 fractions were collected within a 20-42 time window using an automated fraction collector. The chromatograms were monitored at 280 nm.

### Trypsin digestion of SEC fractions

We used bottom-up LC-MS/MS analysis to determine SEC elution profile serum proteins, isolated sMAC, and commercial sMAC. The fractions were introduced into the digestion buffer containing 100 mM Tris-HCl (pH 8.5), 1% w/v sodium deoxycholate (SDC), 5 mM Tris (2-carboxyethyl) phosphine hydrochloride (TCEP) and 30 mM chloroacetamide (CAA). Proteins were digested overnight with trypsin at an enzyme-to-protein-ratio of 1:100 (w/w) at 37 °C. After, the SDC was precipitated by bringing the sample to 1% trifluoroacetic acid (TFA). The supernatant was collected for subsequent desalting by an Oasis µElution HLB 96-well plate (Waters) positioned on a vacuum manifold. The desalted proteolytic digest was dried with a SpeedVac apparatus and stored at -20°C. Prior to LC-MS/MS analysis, the sample was reconstituted in 2% formic acid (FA).

### LC-MS/MS analysis of isolated sMAC SEC fractions

The digested SEC fractions of isolated and commercial sMAC were analyzed using an Ultimate 3000 system (Thermo Scientific) coupled on-line to an Orbitrap Fusion (Thermo Scientific) controlled by Thermo Scientific Xcalibur software. First, peptides were trapped using a 0.3×5 mm PepMap-100 C18 pre-column (Thermo Scientific) of 5 µm particle size and 100 Å pore size prior to separation on an analytical column (50 cm of length, 75 µm inner diameter; packed in-house with Poroshell 120 EC-C18, 2.7 µm). Trapping of peptides was performed for 1 min in 9% solvent A (0.1 % FA) at a flow rate of 0.03 ml/min. The peptides were subsequently separated by a 55 min gradient as follows: 9-13 % solvent B (0.1% FA in 80% CAN) in 1 min, 13-44 %B in 37 min, 44-99 %B in 3 min, 99 %B for 4 min, 99-9 %B in 1 min and finally 9% B for 8 min. The flow was 300 nl/min. The mass spectrometer was operated in a data-dependent mode. Full-scan MS spectra from 375-1600 Th were acquired in the Orbitrap at a resolution of 60,000 with standard AGC target and auto maximum injection time. Cycle time for MS^2^ fragmentation scans was set to 1 s. Only peptides with charge states 1-6 were fragmented, and dynamic exclusion properties were set to n = 1, for a duration of 10 s. Fragmentation was performed using HCD collision energy of 28 % in the ion trap and acquired in the Orbitrap at a resolution of 15,000 and standard AGC target with an isolation window of 1.4 Th and maximum injection time mode set to auto.

### LC-MS/MS analysis of SEC serum fractions

The 74 digested SEC fractions activated or nonactivated serum were analyzed by LC-MS/MS. Separation of digested protein samples was performed on an Agilent 1290 Infinity HPLC system (Agilent Technologies). Samples were loaded on a 100 µm × 20 mm trap column (in-house packed with ReproSil Pur C18-AQ, 3 µm) (Dr. Maisch GmbH, Ammerbuch-Entringen, Germany) coupled to a 50 µm × 500 mm analytical column (in-house packed with Poroshell 120 EC-C18, 2.7 µm) (Agilent Technologies, Amstelveen). 10 µL of digest from each SEC fraction was used and the amount ∼0.1 µg of peptides was loaded on the LC column. The LC-MS/MS run time was set to 60 min with a 300 nL/min flow rate. Mobile phases A (water/0.1% formic acid) and B (80% ACN/0.1% formic acid) were used for 66 min gradient elution: 13– 44% B for 35 min, and 44–100% B over 8 min. Samples were analyzed on a Thermo Scientific Q Exactive™ HF quadrupole-Orbitrap instrument (Thermo Scientific). Nano-electrospray ionization was achieved using a coated fused silica emitter (New Objective) biased to 2 kV. The mass spectrometer was operated in positive ion mode, and the spectra were acquired in the data-dependent acquisition mode. Full MS scans were acquired with 60,000 resolution (at 200 m/z) and at a scan mass range of 375 to 1600 m/z. The AGC target was set to 3 × 10^6^ with a maximum injection time of 20 ms. Data-dependent MS/MS (dd-MS/MS) scan was acquired at 30,000 resolution (at 200 m/z) and with a mass range of 200 to 2000 m/z. AGC target was set to 1 × 10^5^ with a maximum injection time defined at 50 ms. One µscan was acquired in both full MS and dd-MS/MS scans. The data-dependent method was set to isolation and fragmentation of the 12 most intense peaks defined in a full MS scan. Parameters for isolation/fragmentation of selected ion peaks were set as follows: isolation width = 1.4 Th, HCD normalized collision energy (NCE) = 27%.

### LC-MS/MS data analysis

The LC-MS/MS data were searched against UniProtKB/Swiss-Prot human proteome sequence database with MaxQuant software (version 1.5.3.30 or 2.0.3.0). For label-free quantification iBAQ values were selected as output. For profiling of sMAC components in serum, each fraction’s iBAQ values were extracted and normalized to the highest intensity.

### Data analysis and statistical testing

Unless stated otherwise, graphs are comprised of at least three biological replicates. Statistical analyses were performed in GraphPad Prism 8 and are further specified in the figure legends.

## Acknowledgements

The authors would like to acknowledge Wioleta Zelek and Paul Morgan for providing antibodies for the sMAC ELISAs, Piet Aerts for help with the Silver staining, Benjamin Sellner and Urs Jenal for providing the CGSC7740 wildtype and *wbbL+* strain and Remy Muts for critical reading of the manuscript.

## Author contributions

D.J.D., S.H.M. and B.W.B. conceived the project. D.J.D., M.F.L.W. and M.R. performed all bacterial killing assays in serum and supernatant harvesting. D.J.D., M.F.L.W. and M.R. performed flow cytometry. D.J.D., M.F.L.W. and M.R. performed all ELISA, SDS-PAGE or Western blot analysis. M.F.L.W. and M.R. performed bystander erythrocyte lysis assaysD.J.D. performed LPS O-Ag Silver staining. D.J.D. and M.F.L.W. performed the sMAC isolation via HisTrap beads, subsequent SEC and BN-PAGE. M.V.L. and V.F. performed all protein digestions and LC-MS/MS experiments. D.J.D., M.V.L., M.F.L.W. and V.F. analyzed the data. D.J.D., S.H.M. and B.W.B. wrote the manuscript. M.V.L., V.F. and A.J.R.H. proofread the manuscript before submission. All authors approved the final version of the manuscript.

## Competing interests

The authors declare no competing interests.

## Financial disclosure statement

This work was funded by an ERC Starting grant (639209-ComBact, to S.H.M.R), the Utrecht University Molecular immunology HUB (eSTIMATE), the Aspasia grant (Dutch Research Council NWO, to S.H.M.R.), the Netherlands Organization for Scientific Research (NWO) funding the Netherlands Proteomics Centre through the X-omics Road Map program (project 184.034.019, to A.J.R.H.) and fellowship support from the Independent Research Fund Denmark (Project 9036-00007B, to M.V.L.). The funders had no role in study design, data collection and analysis, decision to published or preparation of the manuscript.

## Notes

### Competing Interest Statement

The authors have declared no competing interest.

## References

1. Gasque P. Complement: A unique innate immune sensor for danger signals. Mol Immunol. 2004;41(11 SPEC. ISS.):1089–98.

2. Ricklin D, Hajishengallis G, Yang K, Lambris JD. Complement: A key system for immune surveillance and homeostasis. Nat Immunol. 2010;11(9):785–97.

3. Merle NS, Church SE, Fremeaux-Bacchi V, Roumenina LT. Complement system part I - molecular mechanisms of activation and regulation. Front Immunol. 2015;6(JUN):1–30.

4. Bhakdi S, Tranum-Jensen J. Molecular nature of the complement lesion. Proc Natl Acad Sci U S A. 1978;75(11):5655–9.

5. Müller-Eberhard HJ. The membrane attack complex of complement. Annu Rev Immunol. 1986;4:503–28.

6. Menny A, Serna M, Boyd CM, Gardner S, Praveen Joseph A, Paul Morgan B, et al. CryoEM reveals how the complement membrane attack complex ruptures lipid bilayers. Nat Commun. 2018;(2018):1–11.

7. Doorduijn DJ, Rooijakkers SHM, Heesterbeek DAC. How the Membrane Attack Complex Damages the Bacterial Cell Envelope and Kills Gram-Negative Bacteria. Bioessays. 2019;41(10):e1900074.

8. Joiner KA. Complement evasion by bacteria and parasites. Annu Rev Microbiol. 1988 May;42(4):201–30.

9. Merino S, Camprubi S, Alberti S, Benedi VJ, Tomas JM. Mechanisms of Klebsiella pneumoniae resistance to complement-mediated killing. Infect Immun. 1992;60(6):2529–35.

10. Doorduijn DJ, Rooijakkers SHM, van Schaik W, Bardoel BW. Complement resistance mechanisms of Klebsiella pneumoniae. Immunobiology. 2016 Oct;221(10):1102–9.

11. Abreu AG, Barbosa AS. How Escherichia coli Circumvent Complement-Mediated Killing. Front Immunol. 2017;8(April):1–6.

12. Brown EJ. Interaction of gram-positive microorganisms with complement. Curr Top Microbiol Immunol. 1985;121:159–87.

13. Heesterbeek DAC, Angelier ML, Harrison RA, Rooijakkers SHM. Complement and Bacterial Infections: From Molecular Mechanisms to Therapeutic Applications. J Innate Immun. 2018;1–10.

14. Lewis LA, Ram S. Meningococcal disease and the complement system. Virulence. 2014;5(1):98–126.

15. Barnum SR, Bubeck D, Schein TN. Soluble Membrane Attack Complex : Biochemistry and Immunobiology. 2020;11(November):1–14.

16. Lin RY, Astiz ME, Saxon JC, Saha DC, Rackow EC. Alterations in C3, C4, factor B, and related metabolites in septic shock. Clin Immunol Immunopathol. 1993 Nov;69(2):136–42.

17. Mook-Kanamori BB, Brouwer MC, Geldhoff M, Ende A van der, van de Beek D. Cerebrospinal fluid complement activation in patients with pneumococcal and meningococcal meningitis. J Infect. 2014 Jun;68(6):542–7.

18. Westra D, Volokhina EB, van der Molen RG, van der Velden TJAM, Jeronimus-Klaasen A, Goertz J, et al. Serological and genetic complement alterations in infection-induced and complement-mediated hemolytic uremic syndrome. Pediatr Nephrol. 2017;32(2):297–309.

19. Morgan BP, Walters D, Serna M, Bubeck D. Terminal complexes of the complement system: new structural insights and their relevance to function. Immunol Rev. 2016;274(1):141–51.

20. Doorduijn DJ, Heesterbeek DAC, Ruyken M, de Haas CJC, Stapels DAC, Aerts PC, et al. Polymerization of C9 enhances bacterial cell envelope damage and killing by membrane attack complex pores. PLoS Pathog. 2021 Nov;17(11):e1010051.

21. Mollnes TE, Lea T, Frøland SS, Harboe M. Quantification of the terminal complement complex in human plasma by an enzyme-linked immunosorbent assay based on monoclonal antibodies against a neoantigen of the complex. Scand J Immunol. 1985 Aug;22(2):197–202.

22. Vogel CW, Fritzinger DC. Cobra venom factor: Structure, function, and humanization for therapeutic complement depletion. Toxicon. 2010;56(7):1198–222.

23. Grossman N, Schmetz MA, Foulds J, Klima EN, Jimenez-Lucho VE, Leive LL, et al. Lipopolysaccharide size and distribution determine serum resistance in Salmonella montevideo. J Bacteriol. 1987;169(2):856–63.

24. Preissner KT, Podack ER, Muller-Eberhard HJ. The Membrane Attack Complex of complement: relation of C7 to the metastable membrane binding site of the intermediate complex C5b-7. Vol. 135, Journal of Immunology. 1985.

25. Bayly-Jones C, Bubeck D, Dunstone MA. The mystery behind membrane insertion: A review of the complement membrane attack complex. Philos Trans R Soc B Biol Sci. 2017;372(1726).

26. Heesterbeek DA, Bardoel BW, Parsons ES, Bennett I, Ruyken M, Doorduijn DJ, et al. Bacterial killing by complement requires membrane attack complex formation via surface-bound C5 convertases. EMBO J. 2019 Feb 15;38(4):1–17.

27. Doorduijn DJ, Bardoel BW, Heesterbeek DAC, Ruyken M, Benn G, Parsons ES, et al. Bacterial killing by complement requires direct anchoring of membrane attack complex precursor C5b-7. PLoS Pathog. 2020;16(6 June):1–23.

28. Lachmann PJ, Thompson RA. Reactive lysis: the complement-mediated lysis of unsensitized cells. II. The characterization of activated reactor as C56 and the participation of C8 and C9. J Exp Med. 1970;131(4):643–57.

29. Götze O, Muller-eberhard HJ. Lysis of erythrocytes by complement in the absence of antibody. J Exp Med. 1970;132(5):898–915.

30. Cooper NR, Müller-Eberhard HJ. The reaction mechanism of human C5 in immune hemolysis. J Exp Med. 1970;132(4):775–93.

31. Zipfel PF, Skerka C. Complement regulators and inhibitory proteins. Nat Rev Immunol. 2009;9(10):729–40.

32. Schmidt CQ, Lambris JD, Ricklin D. Protection of host cells by complement regulators. Immunol Rev. 2016;274(1):152–71.

33. Menny A, Lukassen M V., Couves EC, Franc V, Heck AJR, Bubeck D. Structural basis of soluble membrane attack complex packaging for clearance. Nat Commun. 2021 Oct 19;12(1):6086.

34. Joiner KA, Hammer CH, Brown EJ, Cole RJ, Frank MM. Studies on the mechanism of bacterial resistance to complement-mediated killing. I. Terminal complement components are deposited and released from Salmonella minnesota S218 without causing bacterial death. J Exp Med. 1982;155(3):797–808.

35. Krukonis ES, Thomson JJ. Complement evasion mechanisms of the systemic pathogens Yersiniae and Salmonellae. FEBS Lett. 2020;594(16):2598–620.

36. Grossman N, Svenson SB, Leive L, Lindberg AA. Salmonella O antigen-specific oligosaccharide-octyl conjugates activate complement via the alternative pathway at different rates depending on the structure of the O antigen. Mol Immunol. 1990;27(9):859–65.

37. Rawal N, Pangburn MK. Formation of High-Affinity C5 Convertases of the Alternative Pathway of Complement. J Immunol. 2001;166(4):2635–42.

38. Berends ETM, Gorham RD, Ruyken M, Soppe JA, Orhan H, Aerts PC, et al. Molecular insights into the surface-specific arrangement of complement C5 convertase enzymes. BMC Biol. 2015;13(1):93.

39. Fearon DT, Austen KF. Properdin: binding to C3b and stabilization of the C3b dependent C3 convertase. J Exp Med. 1975;142(4):856–63.

40. Daha MR, Van Es LA. Stabilization of homologous and heterologous cell-bound amplification convertases, C3bBb, by C3 nephritic factor. Immunology. 1981;43(1):33–8.

41. Alcorlo M, Tortajada A, Rodríguez de Córdoba S, Llorca O. Structural basis for the stabilization of the complement alternative pathway C3 convertase by properdin. Proc Natl Acad Sci U S A. 2013;110(33):13504–9.

42. Pedersen D V, Roumenina L, Jensen RK, Gadeberg TA, Marinozzi C, Picard C, et al. Functional and structural insight into properdin control of complement alternative pathway amplification. EMBO J. 2017;36(8):1084–99.

43. Joiner KA, Hammer CH, Brown EJ, Frank MM. Studies on the mechanism of bacterial resistance to complement-mediated killing. II. C8 and C9 release C5b67 from the surface of Salmonella minnesota S218 because the terminal complex does not insert into the bacterial outer membrane. J Exp Med. 1982 Mar 1;155(3):809–19.

44. Nemerow GR, Yamamoto KI, Lint TF. Restriction of complement-mediated membrane damage by the eighth component of complement: a dual role for C8 in the complement attack sequence. J Immunol. 1979 Sep;123(3):1245–52.

45. Sharp TH, Koster AJ, Gros P. Heterogeneous MAC Initiator and Pore Structures in a Lipid Bilayer by Phase-Plate Cryo-electron Tomography. Cell Rep. 2016;15(1):1–8.

46. Russo TA, Beanan JM, Olson R, MacDonald U, Cope JJ. Capsular polysaccharide and the O-specific antigen impede antibody binding: a potential obstacle for the successful development of an extraintestinal pathogenic Escherichia coli vaccine. Vaccine. 2009 Jan 14;27(3):388–95.

47. Geelen SPM, Aerts PC, Verhoef J, Fleer A, van Dijk H. Functional microassay of complement activation by pneumococci. J Microbiol Methods. 1992;14(4):257–65.

48. Verduin CM, Jansze M, Hol C, Mollnes TE, Verhoef J, Van Dijk H. Differences in complement activation between complement-resistant and complement-sensitive Moraxella (Branhamella) catarrhalis strains occur at the level of membrane attack complex formation. Infect Immun. 1994;62(2):589–95.

49. Willems E, Alkema W, Keizer-Garritsen J, Suppers A, van der Flier M, Philipsen RHLA, et al. Biosynthetic homeostasis and resilience of the complement system in health and infectious disease. EBioMedicine. 2019 Jul;45:303–13.

50. Ståhl A lie, Kristoffersson AC, Olin AI, Olsson ML, Roodhooft AM, Proesmans W, et al. A novel mutation in the complement regulator clusterin in recurrent hemolytic uremic syndrome. Mol Immunol. 2009;46(11–12):2236–43.

51. Van Den Heuvel L, Riesbeck K, El Tahir O, Gracchi V, Kremlitzka M, Morré SA, et al. Genetic predisposition to infection in a case of atypical hemolytic uremic syndrome. J Hum Genet. 2018;63(1):93–6.

52. Singh B, Su Y-C, Riesbeck K. Vitronectin in bacterial pathogenesis: a host protein used in complement escape and cellular invasion. Mol Microbiol. 2010 Nov;78(3):545–60.

53. Riesbeck K. Complement evasion by the human respiratory tract pathogens Haemophilus influenzae and Moraxella catarrhalis. FEBS Lett. 2020 Feb 13;1–12.

54. Hallström T, Blom AM, Zipfel PF, Riesbeck K. Nontypeable Haemophilus influenzae protein E binds vitronectin and is important for serum resistance. J Immunol. 2009 Aug 15;183(4):2593–601.

55. Griffiths NJ, Hill DJ, Borodina E, Sessions RB, Devos NI, Feron CM, et al. Meningococcal surface fibril (Msf) binds to activated vitronectin and inhibits the terminal complement pathway to increase serum resistance. Mol Microbiol. 2011 Dec;82(5):1129–49.

56. Preissner KP, Podack ER, Müller-Eberhard HJ. SC5b-7, SC5b-8 and SC5b-9 complexes of complement: ultrastructure and localization of the S-protein (vitronectin) within the macromolecules. Eur J Immunol. 1989 Jan;19(1):69–75.

57. Berends ETM, Mohan S, Miellet WR, Ruyken M, Rooijakkers SHM. Contribution of the complement Membrane Attack Complex to the bactericidal activity of human serum. Mol Immunol. 2015;65(2):328–35.

58. Nunn M a, Sharma A, Paesen GC, Adamson S, Lissina O, Willis AC, et al. Complement inhibitor of C5 activation from the soft tick Ornithodoros moubata. J Immunol. 2005;174(4):2084–91.

59. Liu D, Reeves PR. Escherichia coli K12 regains its O antigen. Microbiology. 1994 Jan;140 (Pt 1(1):49–57.

60. Zelek WM, Morgan BP. Monoclonal Antibodies Capable of Inhibiting Complement Downstream of C5 in Multiple Species. Front Immunol. 2021;11(December):612402.

