## Supplementary information for "Soluble MAC is primarily released from MAC-resistant bacteria that potently convert complement component C5"

### S1 – Complement-dependency of bacterial killing and C3a release in serum for *E. coli* strains

a) MAC-sensitive *E. coli* strains (5x$10^8$ bacteria/ml) were incubated in 5% pooled human serum supplemented with C5 conversion inhibitors 6 µg/ml OmCI and 6 µg/ml eculizumab. Bacterial viability was determined after 60 minutes by counting colony forming units (CFU's) and calculating the survival compared to t=0. The horizontal dotted line represents the detection limit of the assay. b) MAC-sensitive (MAC-sens, orange) and MAC-resistant (MAC-res, blue) *E. coli* strains (5x$10^7$ bacteria/ml) were incubated in 5% pooled human serum. Supernatant was collected by spinning down bacteria after 60 minutes. C3a in the supernatant was quantified by ELISA. Data represent individual values of three independent experiments with mean +/- SD.

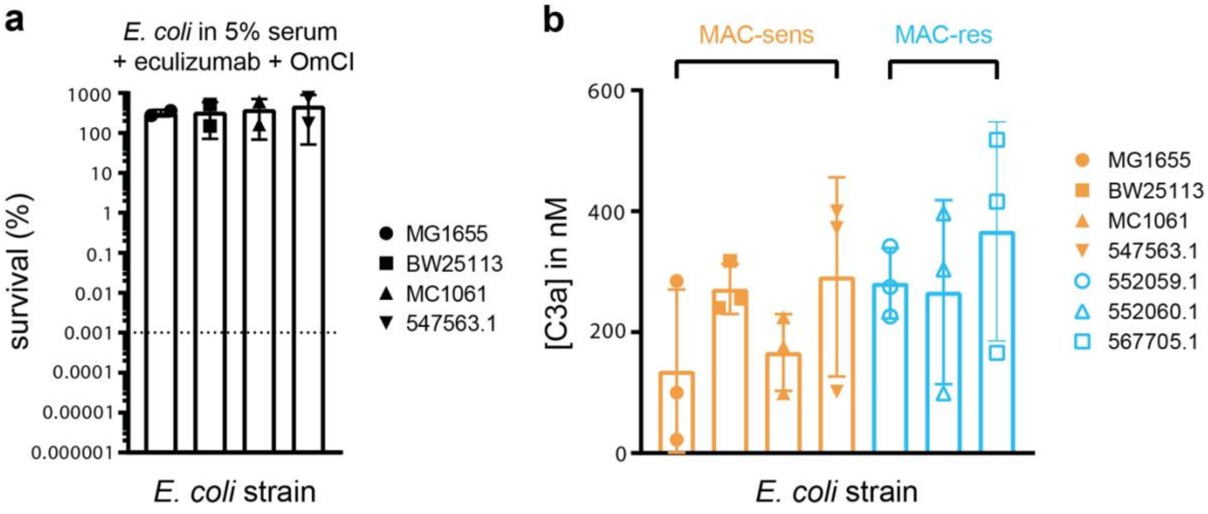

**S2 - Validation specificity sMAC ELISA and sMAC release by *Klebsiella* strains**

sMAC was detected by ELISA in 5% C6-depleted serum supplemented with C6-biotin incubated alone, with 10 µg/ml CVF or 25 µg/ml OmCI for 60 minutes. b) *E. coli* strains (5x10$^{8}$ bacteria/ml) were incubated in 5% C6-depleted serum with C6-biotin, with and without C5 conversion inhibitors OmCI (6 µg/ml) and eculizumab (6 µg/ml). Supernatant was collected by spinning down bacteria after 60 minutes. sMAC was detected in the reaction supernatant (100-fold diluted) by ELISA. c) *Klebsiella* strains (5x10$^{8}$ bacteria/ml) were also incubated in 5% C6-depleted serum with C6-biotin and supernatant was collected by spinning down bacteria after 60 minutes. sMAC was detected in the reaction supernatant (100-fold diluted) by ELISA. Orange strains are MAC-sensitive (MAC-sens), blue strains MAC-resistant (MAC-res) and black strains complement-resistant (comp-res). Serum without bacteria (serum only) was taken as background control. Data represent individual values of one (a, b) or three (c, d) independent experiments with mean +/- SD.

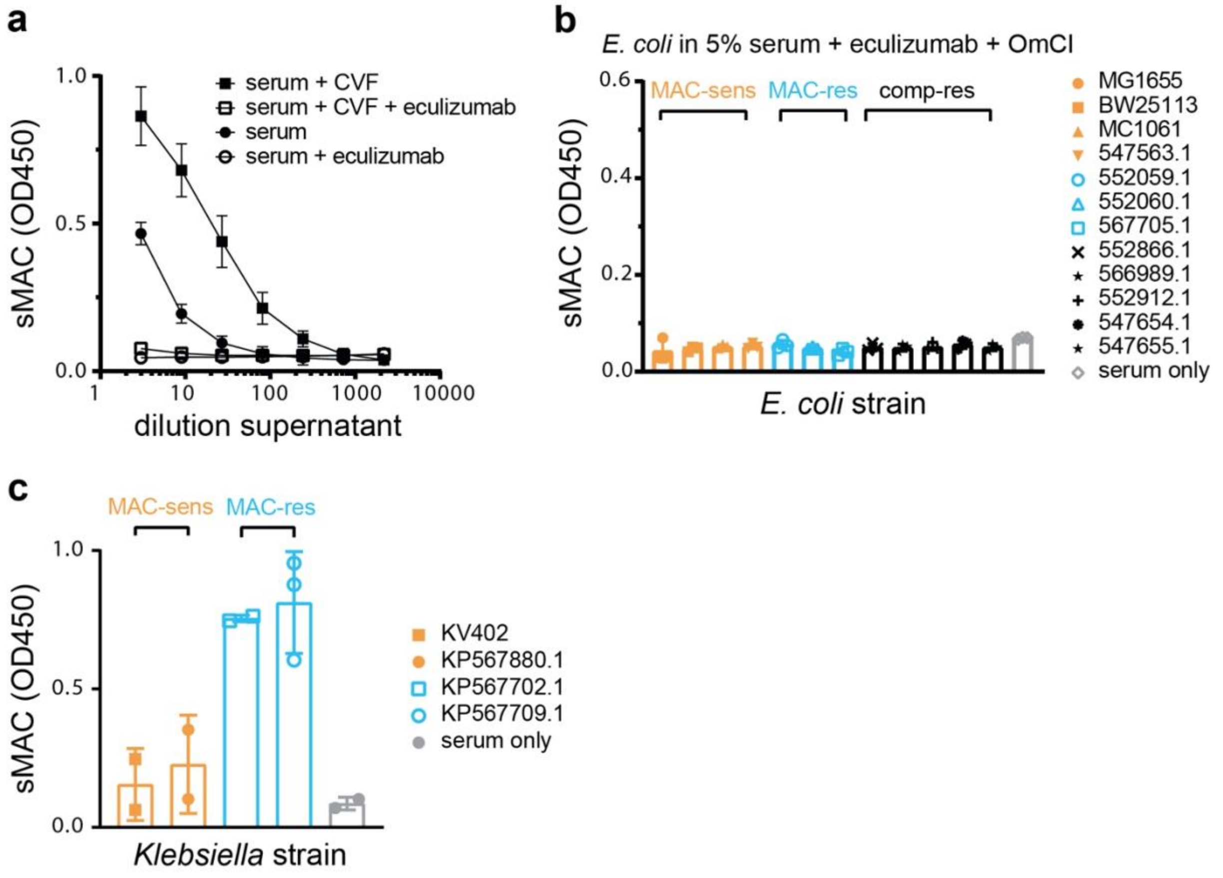

**S3 –LPS O-Ag expression of Gram-negative strains and its effect on C3b deposition**

*E. coli* strains (a) and *Klebsiella* strains (b) were typed for the presence of LPS O-Ag via Silver staining (methods described in [20]). LPS-core were distinguished from LPS O-Ag based on size. Orange strains are MAC-sensitive (MAC-sens), blue strains MAC-resistant (MAC-res) and black strains complement-resistant (comp-res). The Silver stain was previously shown in part (only for Silver MAC-resistant and MAC-sensitive *E. coli* strains) in Doorduijn *et al*. [20]. c) $5x10^7$ bacteria/ml of *E. coli* strain CGSC7740 wildtype (O-Ag$^{neg}$) and *wbbL*+ (O-Ag$^{pos}$) were incubated in a titration of C5-depleted serum for 30 minutes. Bacteria were washed and stained with AF488-labelled mouse monoclonal anti-C3b for 30 minutes. Antibody staining was measured by flow cytometry. Flow cytometry data are represented by individual geoMFI values of the bacterial population. Data represent individual values of two independent experiments with mean +/- SD.

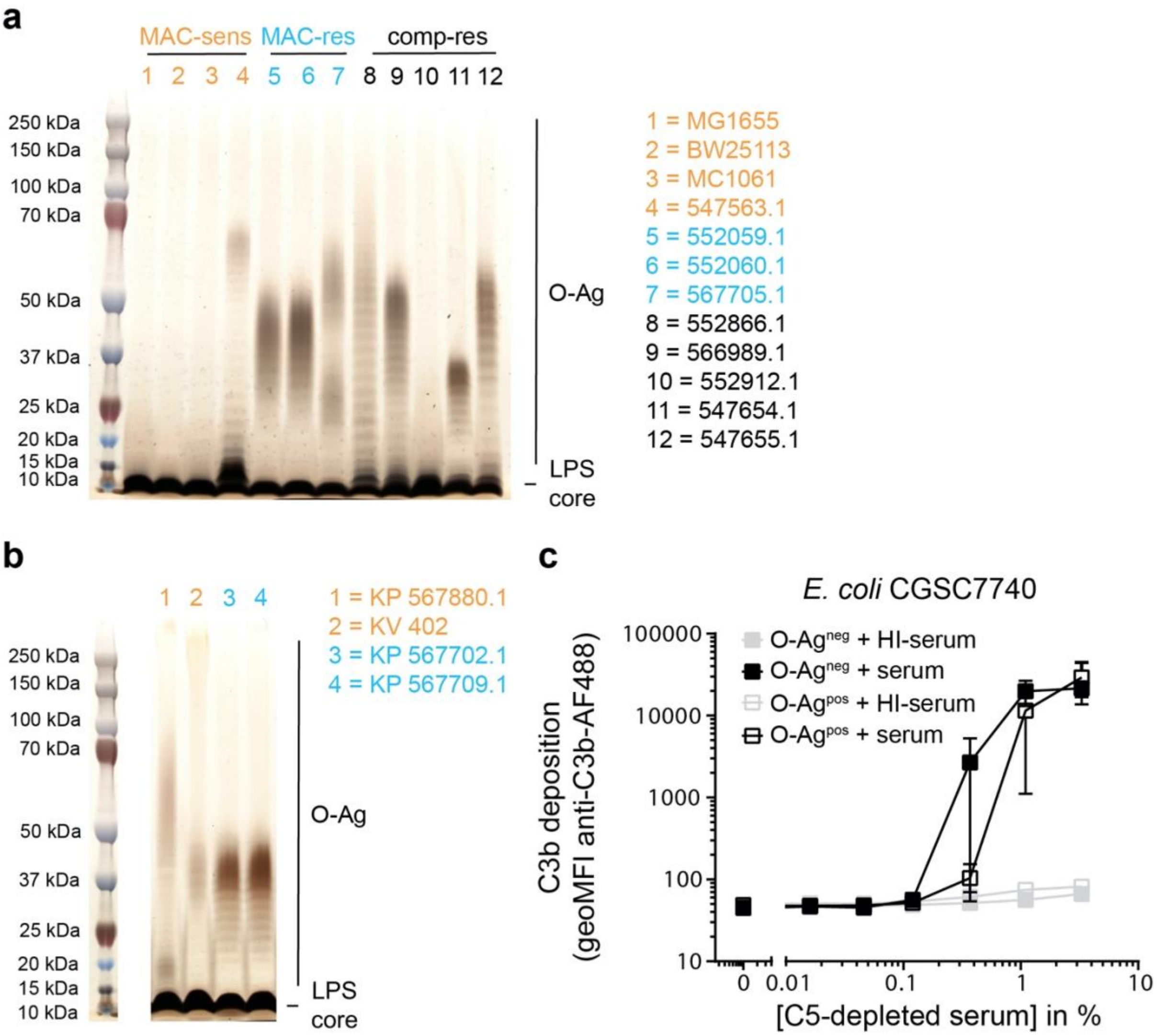

**S4 - sMAC release with purified MAC proteins from convertase-labelled *E. coli* strains.**

MAC-sensitive (MAC-sens, in orange) and MAC-resistant (MAC-res, in blue) *E. coli* strains were labelled with convertases in 10% C5-depleted serum and washed. Next, convertase-labelled bacteria ($3.3x10^8$ per ml) were incubated with alternative pathway (AP) convertase components (50 nM and 20 nM FD) and MAC components (100 nM C5, 100 nM C6, 100 nM C7, 100 nM C8 and 500 nM C9). The supernatant was collected after 60 minutes by spinning down bacteria. sMAC was detected in the reaction supernatant (100-fold diluted) by ELISA. MAC components without bacteria (MAC components only) were taken as background control. Data represent individual values of three independent experiments with mean +/- SD.

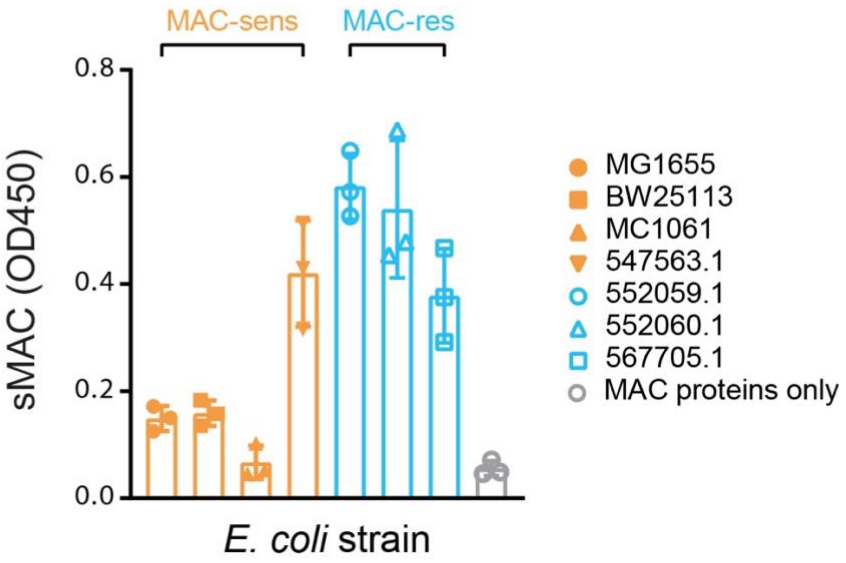

**S5 - Effect of Vn and Clu on C9 polymerization and target-specific MAC assembly.**

MAC-resistant *E. coli* 552059.1 was labelled with convertases in 10% C5-depleted serum and washed. Next, convertase-labelled bacteria (3.3x$10^8$ per ml) were incubated with: alternative pathway (AP) convertase components (50 nM FB and 20 nM FD), MAC components C5-C9 (30 nM C5, 30 nM C6, 30 nM C7, 30 nM C8 and 300 nM Cy5-labelled C9) with or without 133 nM vitronectin (Vn) or 133 nM clusterin (Clu). SDS-PAGE was performed to distinguish monomeric-C9 from polymeric-C9 by in-gel Cy5 fluorescence. b, c) MAC-sensitive *E. coli* MG1655 were labelled with convertases in 10% C5-depleted serum and washed. Next, convertase-labelled bacteria (5x$10^7$ per ml) were incubated with; inner membrane (IM) damage marker 2.5 µM Sytox blue [26], a concentration range of MAC components (C5:C6:C7:C8:C9 as 1:1:1:1:3) alone and with 100 nM Vn or Clu. After 30 minutes, bacteria were analyzed by flow cytometry for binding of Cy5-labelled C9 (b) and Sytox fluorescence as read-out for IM damage (c). Flow cytometry data are represented by geoMFI values of the bacterial population, in the case of Sytox the relative geoMFI was calculated compared to bacteria in buffer and Sytox only. SDS-PAGE images are representative for at least three independent experiments. Data represent mean +/- SD of three independent experiments.

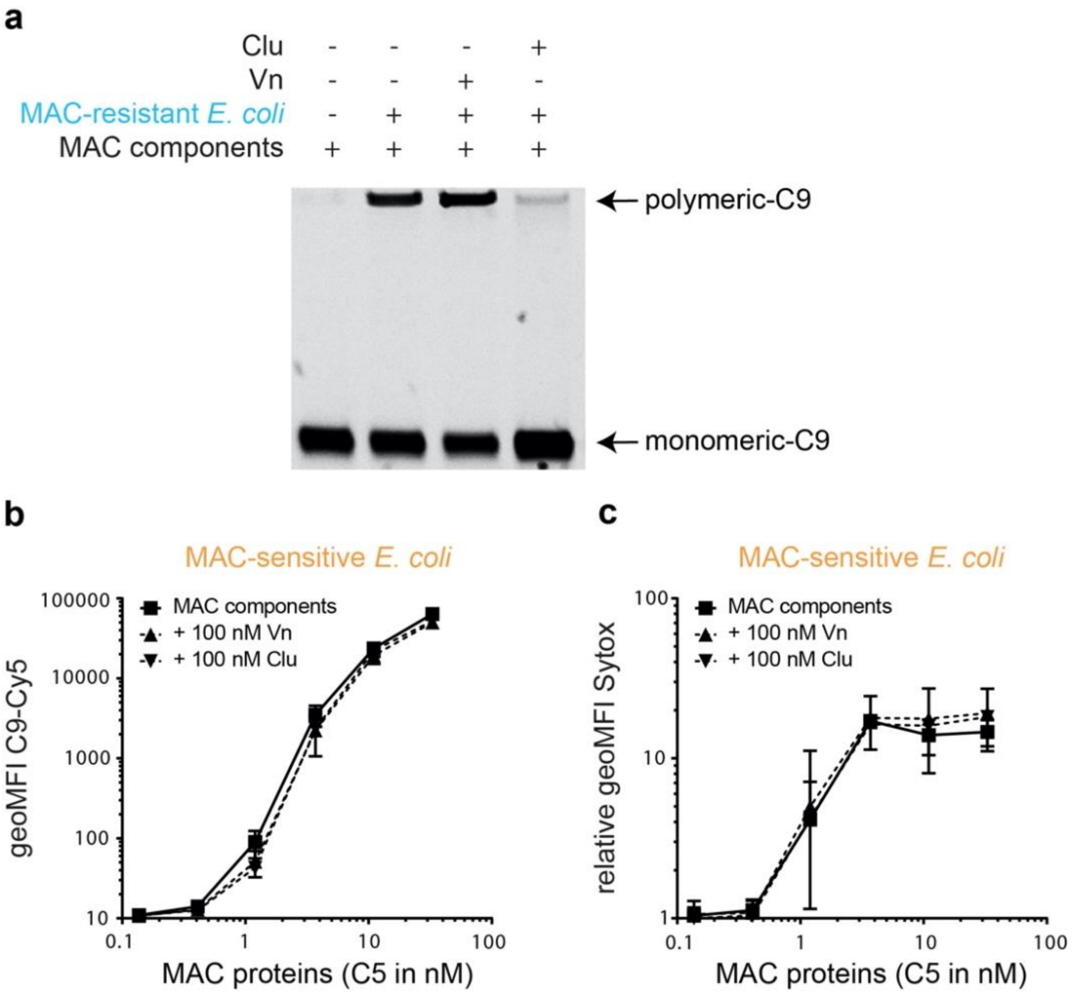

### S6 – Validation isolation sMAC with HisTrap beads and LC-MS/MS SEC fractions

sMAC was detected by ELISA at different steps during the isolation procedure using HisTrap beads. His-tagged sMAC was generated by incubating *E. coli* strain 552059.1 ($5x10^8$ per ml) in 10% C6-depleted serum with 50 nM His-tagged C6 (His-C6) for 60 minutes at 37°C. Supernatant was collected by centrifugation (serum supernatant) and His-tagged sMAC was captured with HisTrap beads for 90 minutes at 4°C. Supernatant was collected from the beads (supernatant HisTrap beads) and sMAC was eluted from beads with 250 mM imidazole (eluate HisTrap beads) for 30 minutes at 4°C. a) sMAC was detected by ELISA for a dilution range of samples. Data represent mean +/- SD of four independent experiments. b-e) LC-MS/MS was performed on individual SEC fractions of concentrated bead eluate for serum incubated with MAC-res *E. coli* and 50 µg commercially available sMAC. The protein abundance of individual sMAC components (iBAQ value) and ratio of individual sMAC components (compared to C5) were represented in for serum incubated with *E. coli* (b, d) and commercial sMAC (c, e). The dotted line (d and e) represents a ratio of 1. LC-MS/MS data represent three individual digests of the same fraction with mean +/- SD that are representative for two independent experiments.

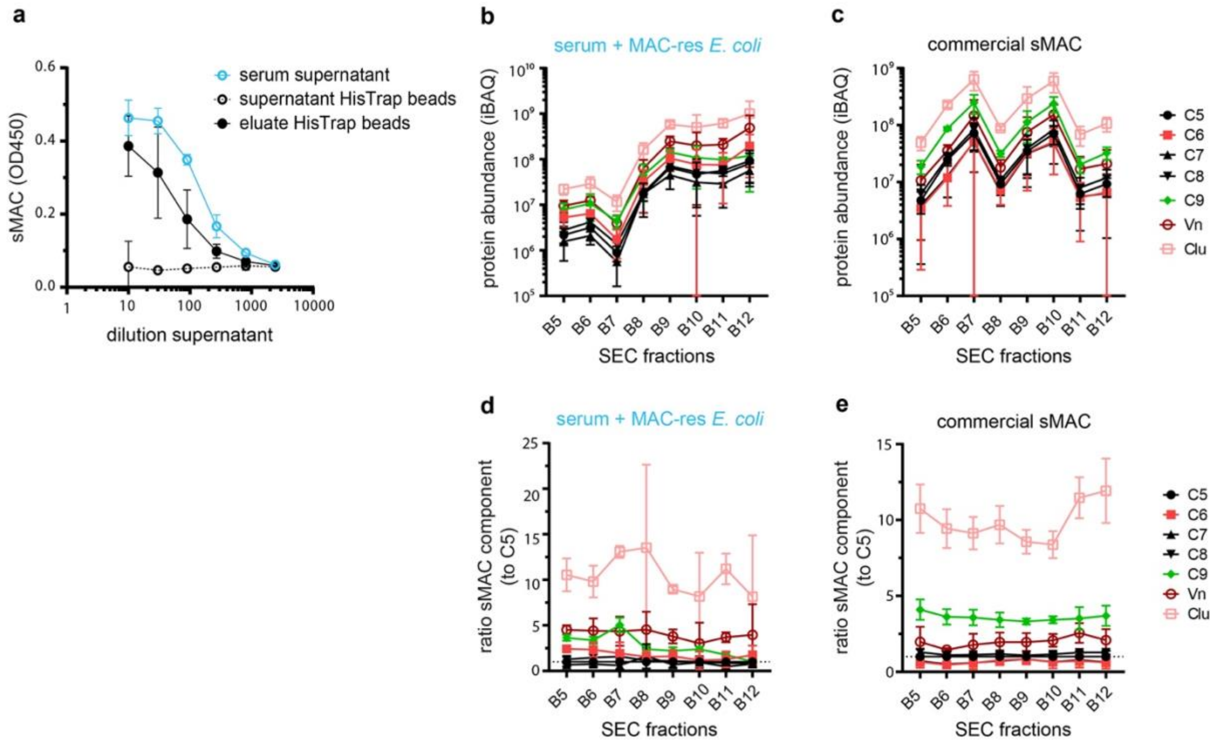
